# Deep Learning Enables Design of Multifunctional Synthetic Human Gut Microbiome Dynamics

**DOI:** 10.1101/2021.09.27.461983

**Authors:** Mayank Baranwal, Ryan L. Clark, Jaron Thompson, Zeyu Sun, Alfred O. Hero, Ophelia Venturelli

## Abstract

Predicting the dynamics and functions of microbiomes constructed from the bottom-up is a key challenge in exploiting them to our benefit. Current ordinary differential equation-based models fail to capture complex behaviors that fall outside of a predetermined ecological theory and do not scale well with increasing community complexity and in considering multiple functions. We develop and apply a long short-term memory (LSTM) framework to advance our understanding of community assembly and health-relevant metabolite production using a synthetic human gut community. A mainstay of deep learning, the LSTM learns a high dimensional data-driven non-linear dynamical system model used to design communities with desired metabolite profiles. We show that the LSTM model can outperform the widely used generalized Lotka-Volterra model. We build methods decipher microbe-microbe and microbe-metabolite interactions from an otherwise black-box model. These methods highlight that Actinobacteria, Firmicutes and Proteobacteria are significant drivers of metabolite production whereas *Bacteroides* shape community dynamics. We use the LSTM model to navigate a large multidimensional functional landscape to identify communities with unique health-relevant metabolite profiles and temporal behaviors. In sum, the accuracy of the LSTM model can be exploited for experimental planning and to guide the design of synthetic microbiomes with target dynamic functions.

## INTRODUCTION

Microbial communities perform chemical and physical transformations to shape the properties of nearly every environment on Earth from driving biogeochemical cycles to mediating human health and disease. These functions performed by microbial communities are shaped by a multitude of abiotic and biotic interactions and vary as a function of space and time. The complex dynamics of microbial communities are influenced by pairwise and higher-order interactions, wherein interactions between pairs of species can be modified by other community members [1, 2, 3]. In addition, the interactions between community members can change as a function of time due to variation in the abiotic environment as well as environmental modification by the microbial community [4]. Therefore, flexible modeling frameworks that can capture the complex and temporally changing interactions that determine the dynamic behaviors of microbiomes are needed. These predictive modeling frameworks could be used to guide the design of precise interventions to manipulate community-level functions to our benefit.

The generalized Lotka-Volterra (gLV) model has been widely used to predict community dynamics and deduce pairwise microbial interactions shaping community assembly [5]. For example, the gLV model has been used to predict the assembly of tens of species based on absolute abundance measurements of lower species richness (i.e. number of species) communities [6, 7, 8]. The parameters of the gLV model can be efficiently inferred based on properly collected absolute abundance measurements and can provide insight into significant microbial interactions shaping community assembly. However, the model does not represent higher-order interactions or microbial community functions beyond species growth. To capture such microbial community functions, hybrid gLV models have been developed to predict a community-level functional activity based on species abundance [8, 9]. However, these approaches have been limited to the prediction of a single community-level function at a single time point. Therefore, new modeling frameworks are needed to capture temporal changes in multiple community-level functions, such as tailoring the metabolite profile of the human gut microbiome [10].

Deep machine learning approaches, such as recurrent neural networks (RNNs), are universal function approximators [11, 12] that enable greater flexibility compared to gLV models for modeling dynamical systems. However, deep learning models often require significantly more model parameters, which poses additional challenges to model fitting and generalizability. A particular RNN model architecture called long short-term memory (LSTM) addresses challenges associated with training on sequential data by incorporating gating mechanisms that learn to regulate the influence of information from previous instances in the sequence [13]. From their initial successes in speech recognition [14] and computer vision [15], LSTMs have recently been applied to modeling biological data such as subcellular localization of proteins [16] and prediction of biological age from activity collected from wearable devices [17]. Related to microbiomes, deep learning frameworks have been applied to predict gut microbiome metabolites based on community composition data [18], final community composition based on microbial interactions [19] and end-point community composition based on the presence/absence of species [20]. In addition, RNN architectures have been used to model phytoplankton [21] and macroinvertebrate [22] community dynamics. Despite achieving reasonable prediction performance, previous efforts at modeling ecological system dynamics using RNNs are typically limited to handful of organisms (<10), have provided limited model interpretation and have not been leveraged to predict temporal changes in community behaviors. In addition, RNN architectures have not been used for bottom-up community design, which could be exploited for applications in bioremediation, bioprocessing, agriculture and human health [23, 24, 25].

Here we apply LSTMs to model time dependent changes in species abundance and production of key health-relevant metabolites of a diverse 25-member synthetic human gut community. We use the trained model to elucidate significant microbe-microbe and microbe-metabolite interactions. The flexibility and accuracy of the LSTM model enabled systematic integration into our experimental planning process, in two stages. First the LSTM was fit to an initial pilot experiment with low temporal resolution involving a moderate number of synthetic microbial communities. These communities were selected uniformly at random from the tens of millions of possible communities that could be experimentally explored. The distribution of LSTM metabolite predictions was then used to identify sparse sub-communities in the tails of the distribution, communities that we refer to as “corner cases”. A second experiment was then performed that expands the training data for the LSTM in the vicinity of these corner cases with higher time resolution. The LSTM-guided two-stage experimental planning procedure substantially reduced the number of experiments compared to random sampling of the functional landscape with temporal resolution in a single stage experiment. Therefore, the LSTM analysis enabled our main findings on dynamical behaviors of communities and identified the key species critical for growth and that shape metabolite profiles. Compared to the gLV model, the proposed LSTM framework provides a better fit to the experimental data, captures higher-order interactions and provides higher accuracy predictions of species abundance and metabolite production. In addition, our approach preserves model interpretability through a suitably developed gradient-based framework and locally interpretable model-agnostic explanations (LIME) [26]. Using our time-series data of species abundance and metabolite concentrations, we demonstrate that the temporal behaviors of the communities cluster into distinct groups based on the presence and absence of sets of species. Our results highlight that LSTM models are powerful tools for predicting and designing the dynamic behaviors of microbial communities.

## RESULTS

### LSTM accurately predicts microbial community assembly

Our first objective was to determine if the LSTM model could capture the temporal changes in species abundance in response to dilution, which results in changes in nutrient availability and mortality [27]. We tested the effectiveness of the proposed LSTM method on the time-resolved species abundance data of a well-characterized twelve-member synthetic human gut community [6]. The experimental data consists of species abundance sampled approximately every 12 hours. A total of 175 microbial communities with sizes varying from 2 to 12 were used to train and evaluate the proposed LSTM model. To represent temporal variation in cell densities and nutrient availability, the community was diluted by 20-fold every 24 hours into fresh media (**Fig. 1a**). The dilution of the community introduces further complexity towards model training with external perturbations to the competitive environment. The experimental data was split into non-overlapping training and hold-out test sets, and an appropriate LSTM network was trained to predict species abundances at various time points given the information of initial species abundance. The details on the train/test split and the number of model hyperparameters are provided in Table S2. We found that a total of five LSTM units can predict species abundance at different time points (12, 24, 36, 48 and 60 hours) based on the initial species abundance. The output of each LSTM unit is used as an input to the next unit. However, the input to the current LSTM unit is randomized between the output from the previous LSTM unit and the true abundance at the current time point in the randomized teacher forcing mode of training in order to eliminate temporal bias in the prediction of end-point abundances. We did not model the effect of dilution explicitly, since the experimental procedure was consistent across all communities. This also highlights the advantage of using black-box approaches, such as the LSTM network, where physical parameters such as dilution do not need to be explicitly modeled.

**Figure 1:**
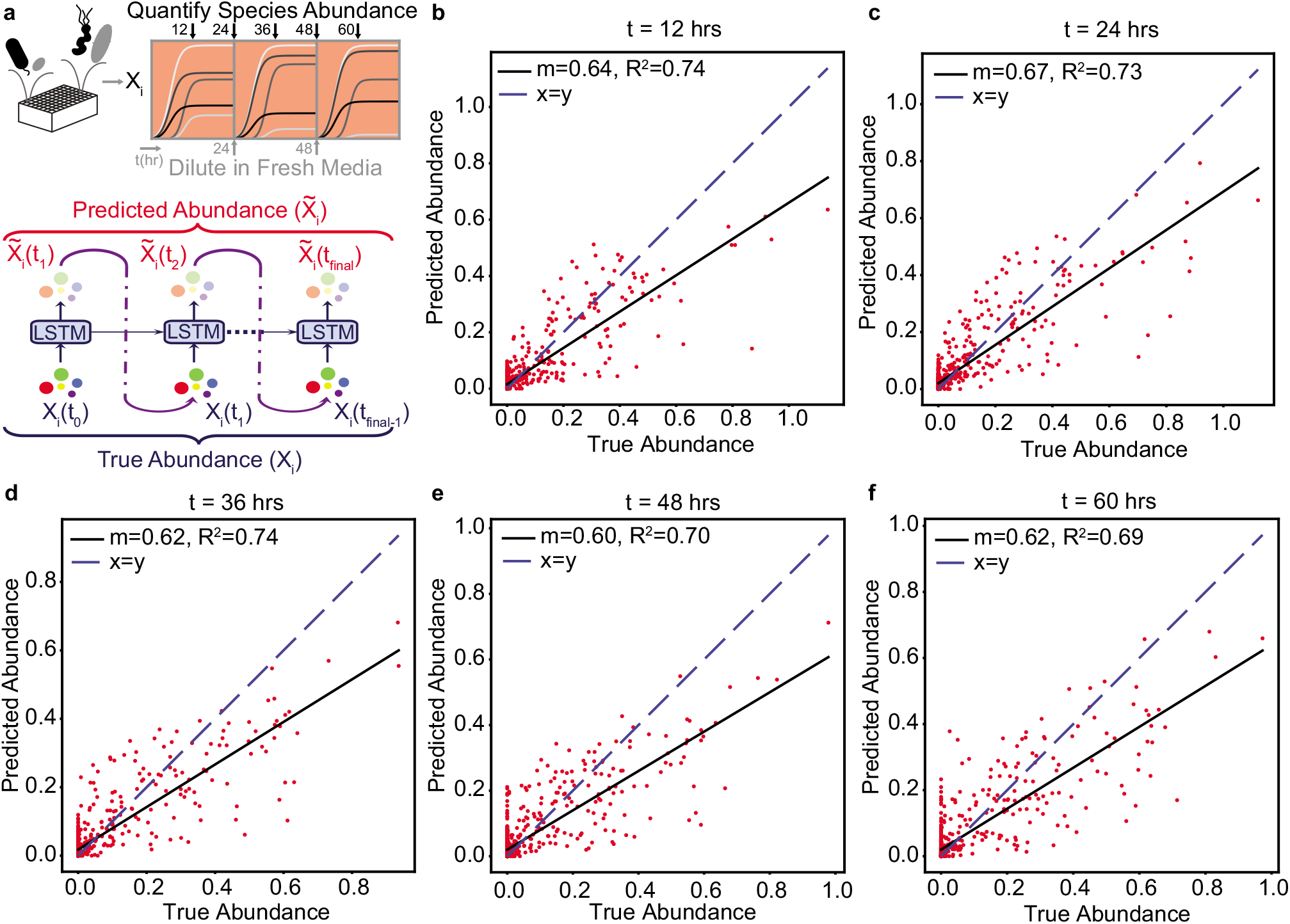
LSTM model can predict the temporal changes in species abundance in a 12-member synthetic human gut community in response to dilution. **(a)** Proposed LSTM modeling methodology for the dynamic prediction of species abundance in a microbial community. The initial abundance information is an input to the first LSTM cell, the output of which is trained to predict abundance at the next time point. Consequently, the predicted abundance becomes an input to another LSTM cell with shared weights to predict the abundance at the subsequent time point. The process is repeated until measurements at all time points are available. **(b)** Scatter plot of measured (true) and predicted species abundance of a 12-member synthetic human gut community at 12 hr (*N* = 876, p-value = 2.44*e* − 257). **(c)** Scatter plot of measured (true) and predicted abundance at 24 hr (p-value = 6.51*e* − 257). **(d)** Scatter plot of measured (true) and predicted abundance at 36 hr (p-value = 7.42*e* − 257). **(e)** Scatter plot of measured (true) and predicted abundance at 48 hr (p-value = 1.66*e* − 227). **(f)** Scatter plot of measured (true) and predicted abundance at 60 hr (p-value = 3.39*e* − 227).

Despite the periodic change in dilution and variations in the sampling times, the proposed LSTM method accurately predicts (Pearson *R*^2^-scores of 0.74, 0.73, 0.74, 0.70 and 0.69 at time points 12, 24, 36, 48 and 60 hours, respectively) not only the end-point species abundance, but also the abundances at intermediate time points on hold-out test sets (**Fig. 1b-1f**). These results demonstrate that the LSTM model can accurately predict the temporal changes in species abundance of multi-species communities in the presence of external perturbations.

### LSTM outperforms the generalized Lotka Volterra ecological model

The gLV model is a widely used ecological model consisting of a coupled set of ordinary differential equations that captures the growth dynamics of members of a community based on their intrinsic growth rate and interactions with all pairs of constituent community members [6]. Therefore, gLV models are not suited to capture higher-order interactions among species or changes in inter-species interactions resulting from variation in the environment. By contrast, the LSTM modeling framework is flexible and can capture complex relationships between species as well as time-dependent changes in inter-species interactions. To quantify these differences, we characterized the performance of the gLV and LSTM models in response to third-order interaction perturbations that varied in magnitude to evaluate the strengths and limitations of these modeling frameworks.

We consider a gLV model of a 25-member microbial community whose dynamics are governed by single organism growth and whose pairwise interactions match those inferred in a previous study [25]. Using this model, we simulate sub-communities that vary in the number of species. Of all the randomly simulated communities, those containing six or fewer species are used to train both the gLV and LSTM models (624 training communities), while the remaining communities (3299 test communities with 10 or more species) are used as a hold-out test set. The simulated data spans 48 hours separated by an interval of 8 hours, reflecting the experimentally feasible periodic sampling interval of 8 hours.

The performances of the trained gLV and LSTM models on the hold-out test sets are similar and are able to accurately predict the trends in species abundance (Pearson *R*^2^ of 0.89 and 0.85 for gLV and LSTM models, respectively) (**Fig. 2b,c** left). Since the training and test data is based on the gLV model, the performance of the gLV is moderately better than the LSTM model. We next explore the scenario where the simulated model comprises low magnitude (mild) third-order interactions (third-order interaction coefficients that do not exceed 25% of the maximum of the absolute values of the coefficients for the second-order interactions). In this case, the performance of the LSTM model is substantially better than the gLV model with the *R*^2^-score of 0.85, as opposed to 0.52 for the gLV model (**Fig. 2b,c**, middle). In addition, the LSTM model performs significantly better than the gLV model for higher magnitude (moderate) third-order perturbations (third-order interaction coefficients that do not exceed 50% of the maximum of the absolute values of the coefficients for second-order interactions) (**Fig. 2b,c**, right).

**Figure 2:**
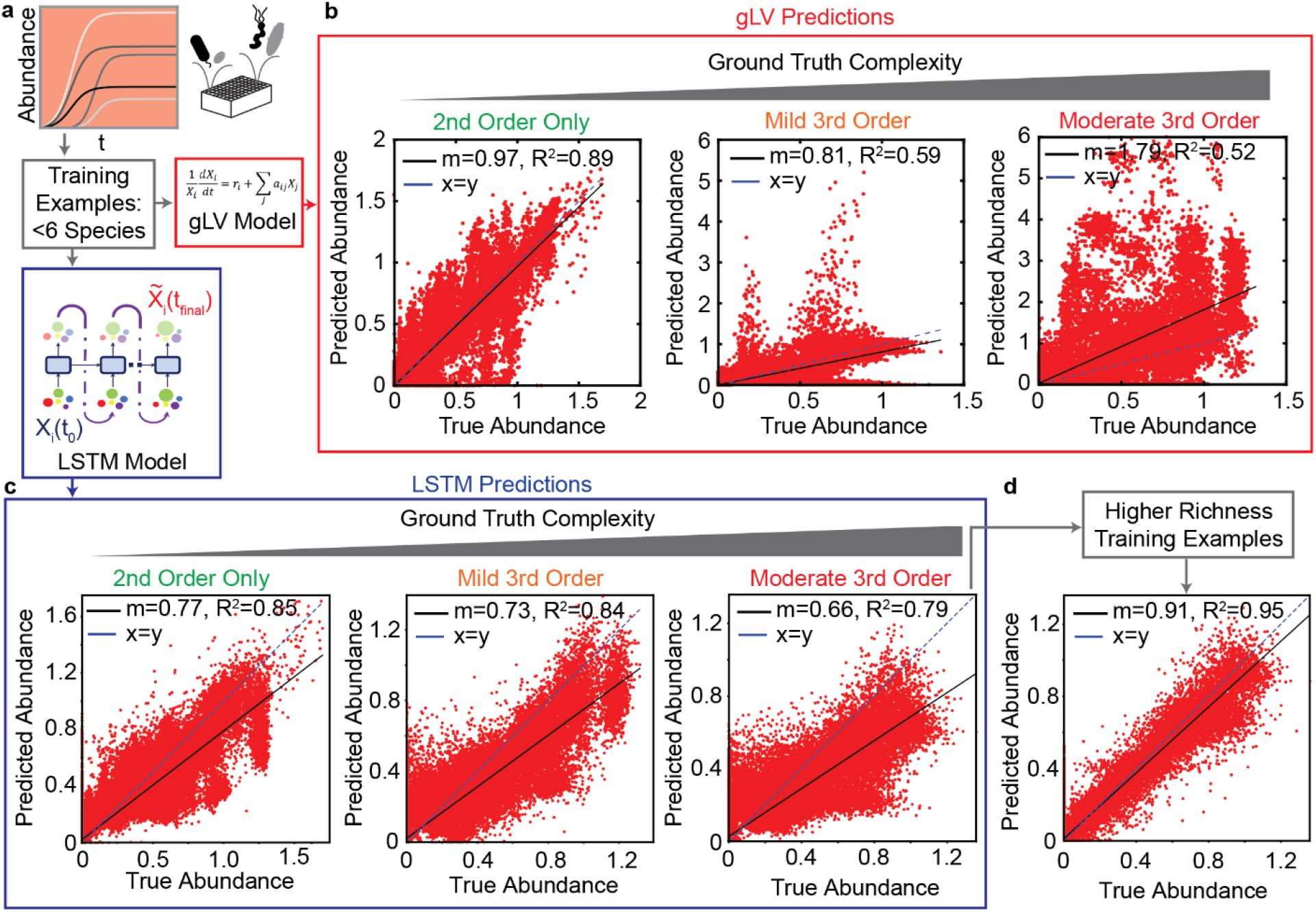
Comparison of generalized Lotka Volterra (gLV) and Long Short Term Memory (LSTM) model prediction performance of species abundance in a 25-member microbial community in response to third-order perturbations of varying magnitude. For both models, training data consists of low species richness communities (≤ 6 species, *N* = 82,475, p-value < 0.0001). **(a)** & **(d)**: Data was generated using a gLV model that captures single species growth and pairwise interactions. Scatter plots of true versus predicted species abundance at *t* = 48hr using the gLV and LSTM models, respectively. **(b)** & **(e)** Scatter plot of true versus predicted species abundance of the gLV and LSTM models, respectively when the simulated data is subjected to low magnitude (mild) third-order interactions. **(c)** & **(f)** Scatter plot of true versus predicted species abundance of gLV and LSTM models, respectively when the simulated data is further subjected to moderately large third order interactions. **(g)** Scatter plot of true versus predicted species abundance for the LSTM model. The training set included a set of higher richness communities (50 each of 11 & 19 member communities).

This in silico analysis reflects the significance of adopting more expressive neural network models over severely constrained parametric models, such as, gLV. In addition, a key advantage of the proposed LSTM model over the gLV model is the amount of time required for training the two models. Note that the gLV equations are coupled nonlinear ordinary differential equations, and thus training gLV models requires substantial computational time (nearly 5-6 hours) whereas the LSTM models can be trained in less than 2 minutes on the same platform. Therefore, the LSTM approach is highly suited for real-time training and planning of experiments. The details on the computational hardware are provided in the Methods section.

We also observed a crescent shaped prediction profile, representing an inherent bias and thus indicating that the species abundances are underpredicted by the LSTM model for the small community training set (**Fig. 2c**). Using the in silico experiments, we aim to not just compare the performances of the gLV and LSTM models, but also to identify what type of datasets are required for building predictive models of high richness community behaviors depending on the nature of their underlying interactions. Thus, we created a new training set consisting of communities containing moderately strong third-order interactions which the gLV model fails to capture. To counter the aforementioned LSTM bias, we augmented the training set with 100 communities enriched with a larger number of species (randomly sampled 11 and 19-member communities). Using this enriched training set, the LSTM network accurately predicts the community dynamics of the hold-out set with an *R*^2^ of 0.95 (**Fig. 2d**). Our results show that the prediction bias is eliminated when the training set includes a set of higher species richness communities. In sum, the LSTM has difficulty predicting the behavior of high richness communities when the training data only consists of low richness communities. However, adding a moderate number of high richness communities to the training set considerably improves the prediction performance of the LSTM.

### LSTM enables end-point design of multifunctional synthetic human gut microbiomes

While predicting the abundance of microbial species is useful, the chemical transformations (i.e. functions) performed by the community are the key design variables for microbiome engineering goals, including benefiting human health [28]. Thus, we were motivated to further explore prediction of microbial community functions using the LSTM framework based on our success in predicting community dynamics combined with the ease of incorporating additional output variables. Therefore, we applied the LSTM framework to design health-relevant metabolite profiles using synthetic human gut communities.

A core function of gut microbiota is to transform complex dietary substrates into fermentation end products such as the beneficial metabolite butyrate, which is a major determinant of gut homeostasis [29]. In a previous study, we designed butyrate-producing synthetic human gut microbiomes from a set of 25 prevalent and diverse human gut bacteria using a hybrid gLV and statistical model. This hybrid model consists of a gLV model for predicting community assembly and a linear regression model with interactions to predict butyrate production from species absolute abundance at a given time point [25]. While the hybrid model approach was successful for predicting butyrate concentration, designing community-level metabolite profiles rather than optimizing the concentration of a single metabolite adds substantial complexity and limited flexibility using the hybrid modeling approach. Thus, we leveraged the accuracy and flexibility of LSTM models to design the metabolite profiles of synthetic human gut microbiomes. We focused on the fermentation products butyrate, acetate, succinate, and lactate which play important roles in the gut microbiome’s impact on host health and interactions with constituent community members [10].

We used the species abundance and metabolite concentrations from our previous work [25] to train an LSTM model. This model uses a feed-forward network (FFN) at the output of the final LSTM unit that maps the endpoint species abundance to the concentrations of the four metabolites (**Fig. 3a**). The entire neural-network model comprising LSTM units and a feed-forward network is learned in an end-to-end manner during the training process, (i.e., all the network weights are trained simultaneously). Cross-validation of this model (Model M1, **Table S1**) on a set of hold-out community observations shows good agreement between the model predictions and experimental measurements for metabolite concentrations and microbial species abundances (**Fig. S1**). Thus, we used this model to design species-rich (i.e. >10 species) microbial communities with tailored metabolite profiles (**Fig. 3a**).

**Figure 3:**
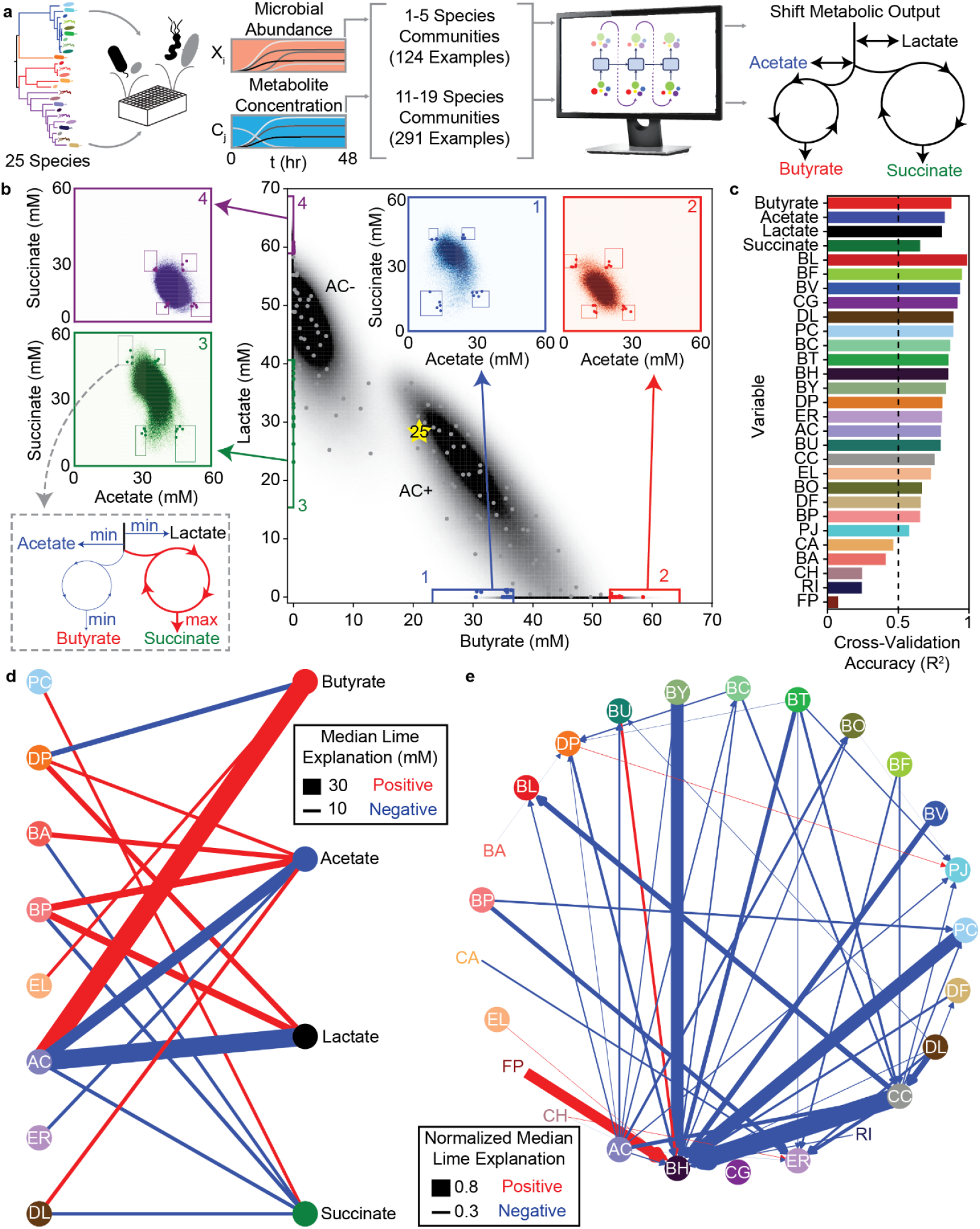
LSTM-guided design and interpretability of community-level metabolite production profiles. **(a)** Schematic of model-training and design of communities with shifted metabolite outputs. **(b)** Heat map of butyrate and lactate concentrations of all possible communities predicted by the LSTM model M1. Grey points indicate communities chosen via *k*-means clustering to span metabolite design space. Colored boxes indicate “corner” regions defined by 95^th^ percentile values on each axis with points of the corresponding color indicating designed communities within that “corner”. Insets show heat maps of acetate and succinate concentrations for all communities within the corresponding boxes on the main figure. Boxes on the inset indicate “corners” defined by 95^th^ percentile values on each axis with colored points corresponding to the same points indicated on the main plot. **(c)** Cross-validation accuracy of LSTM model trained and validated on a random 90/10 split of all community observations (model M2), evaluated as Pearson correlation *R*^2^ for the correlation of predicted versus measured for each variable (all p-values< 0.05, N and p-value for each test reported in **Table S3**). Dashed line indicates *R*^2^ = 0.5, which is used as a cutoff for including a variable in the subsequent network diagrams. **(d)** and **(e)** Network representation of median LIME explanations of the LSTM model M2 from **(c)** for prediction of each metabolite concentration **(d)** or species abundance **(e)** by the presence of each species. Edge widths are proportional to the median LIME explanation across all communities from **(b)** used to train the model in units of concentration (for **(d)**) or normalized to the species’ self-impact (for **(e)**). Only explanations for those variables where the cross-validated predictions had *R*^2^ > 0.5 are shown. Networks were simplified by using lower thresholds for edge width (5 mM for **(d)**, 0.2 for **(e)**). Red and blue edges indicate positive and negative contributions, respectively.

We first used the LSTM model M1 to simulate every possible combination of >10 species (26,434,916 communities). The simulated communities separate into two regions: one centered around a dense ellipse of high butyrate concentration characterized by communities containing the butyrate-producing species *Anaerostipes caccae* (AC) and a second dense ellipse of low butyrate concentration characterized by communities lacking AC (**Fig. 3b**). This bimodality due to the presence/absence of AC is consistent with our previous finding that AC is the strongest driver of butyrate production in this system [25]. In addition, the strong negative correlation between lactate and butyrate in the AC+ ellipse (*R*^2^ = 0.72, *p* < 0.001, N=14,198,086) is consistent with the ability of AC to convert lactate into butyrate [25]. These results demonstrate that the LSTM model can capture the major microbial drivers of metabolite production as well as correlations between different metabolites.

We used our simulated metabolite production landscape to plan informative experiments for testing the capabilities of our model. First, we designed a set of “distributed” communities that spanned the range of typical metabolite concentrations predicted by our model. To this end, we selected 100 communities that fell closest to the centroids of 100 clusters determined using k-means clustering of the 4-dimensional metabolite space. Second, we designed a set of communities to test our model’s ability to predict extreme shifts in metabolite outputs. To do so, we identified four “corners” of the distribution in the lactate and butyrate space (**Fig. 3b**). We next examined the relationship between acetate and succinate within each of these corners and found that the distributions varied depending on the given corner (**Fig. 3b**, inset). The total carbon concentration in the fermentation end products across all predicted communities displayed a narrow distribution (mean 316 mM, standard deviation 20 mM, **Fig. S2**). The production of the four metabolites are coupled due to the structure of the metabolic networks and fundamental stoichiometric constraints [30]. Therefore, the model learned the inherent “trade-off” relationships between these fermentation products based on the patterns in our data. We chose a final set of “corner” communities for experimental validation by choosing 5 communities from each combination of maximizing or minimizing each metabolite (80 communities total, see **Methods** for details).

By experimentally characterizing the 180 designed communities, we found that the LSTM model M1 accurately predicted the rank order of metabolite concentrations and microbial species abundances, substantially outperforming a composite model (gLV and regression) trained on the same data for the majority (59%) of output variables (**Fig. S3a**). Notably, the LSTM model prediction accuracy for the metabolites was similar for both the “distributed” and “corner” communities (**Fig. S3b-e**). These results indicate that our model is useful for designing communities with a broad range of metabolite profiles that includes the extremes of the distributions. To understand how well our model could separate groups of communities with extreme behaviors, we treated the “corners” as classes and quantified the classification accuracy of our model. The model accurately classified the communities when considering only butyrate and lactate concentrations. However, the model had poorer separation when acetate and succinate were also considered in defining the classes (**Fig. S3f**). The misclassification rate was higher for small Euclidean distances between classes and decreased with the Euclidean distance (**Fig. S3g**). This implies that the insufficient variation in concentrations due to fundamental stoichiometric constraints limited our ability to define 16 distinct classes that maximized/minimized each metabolite. While model M1 accurately predicted metabolite concentrations and the majority of species abundances, the predictions of several individual species were still quite poor (*R*^2^ = 0 − 0.6, **Fig. S3a**). Thus, we used the dataset to improve the model. To this end, we combined the new observations with the original observations and randomly partitioned the data into 90% for training and 10% for cross-validation. The resulting model (M2, **Table S1**) was substantially more predictive of species abundances (*R*^2^ > 0.5 for all but five species FP, RI, CA, BA, CH (**Fig. 3c**).

One of the commonly noted limitations of machine learning models is their lack of interpretability for extracting biological information about the system. Thus, we used our predictive LSTM model to decipher key relationships among variables to deepen our biological understanding of the system. We used local interpretable model-agnostic explanations (LIME) [31] to quantify the impact of each species’ presence on the prediction of each metabolite concentration and species abundance in each of the sub-communities used to train model M2. We used the median impact of each species presence on each of the metabolite concentrations and species abundances across all training instances to generate networks that provided key insights into microbe-metabolite (**Fig. 3d**) and microbe-microbe (**Fig. 3e**) interactions. In general, these networks represent broad design principles for community metabolic output by indicating which species have the most consistent and strong impacts on each metabolite and species abundance across a wide range of sub-communities. For instance, the metabolite network highlights AC as having the largest positive effect on butyrate production with additional positive contribution from EL and negative contribution from DP, consistent with the previous hybrid gLV model of butyrate production by this community [25]. Additionally, the number of microbial species impacting each metabolite in these networks trended with the number of microbial species in the system that individually produced or consumed each metabolite (**Fig. S4**). For example, butyrate displayed the fewest edges (3) and was produced by the lowest number of individual species (4). By contrast, acetate had the most edges (6) and was produced by the largest number of individual species (19). The inferred microbe-metabolite network consisted of diverse species including Proteobacteria (DP), Actinobacteria (BA, BP, EL), Firmicutes (AC, ER, DL) and one member of Bacteroidetes (PC) but excluded members of *Bacteroides*. Therefore, while *Bacteroides* exhibited high abundance in many of the communities, they did not substantially impact the measured metabolite profiles but instead modulated species growth and thus community assembly (**Fig. 3e**).

The LIME explanations of inter-species interactions exhibited a statistically significant correlation with their corresponding inter-species interaction parameters from a previously parameterized gLV model of this system [25] (**Fig. S5**). The sign of the interaction was consistent in 80% of the interactions with substantial magnitude (> 0.05 in both the LIME explanations and gLV parameters). This consistency with previous observations suggests that the LSTM model was able to capture the same broad trends in interspecies relationships as gLV (interpreted through the average LIME explanation across all observed communities). The LSTM model captured more nuanced context-specific behaviors (interpreted as the LIME explanation for one specific community context) than the mathematically restricted gLV model, which substantially improved the LSTM model’s predictive capabilities. These results demonstrate that the LSTM framework is useful for developing high accuracy predictive models for the design of precise community-level metabolite profiles. Our approach also preserves the ability to decipher different types of interactions in the LSTM model that are explicitly encoded in less accurate and flexible mechanistic models such as gLV.

### Sensitivity of prediction accuracy highlights poorly understood species and pairwise interactions

Identification of species that limit prediction performance could guide selection of informative experiments to deepen our understanding of the behaviors of poorly predicted communities. Therefore, we evaluated the sensitivity of the LSTM model prediction accuracy to species presence/absence and the amount of training data. High sensitivity of model prediction performance to the number of training communities indicates that collection of additional experimental data would continue to improve the model. Additionally, identifying poorly understood communities will guide ML-informed planning of experiments. To evaluate the model’s sensitivity to the size of the training dataset, we computed the hold-out prediction performance (*R*^2^) as a function of the size of the training set by sub-sampling the data (**Fig. 4a**). We used 20-fold cross-validation to predict metabolite concentrations and species abundance. Our results show that the ability to improve prediction accuracy as a function of the size of the training data set was limited by the variance in species abundance in the training set (**Fig. S6**). For instance, certain species with low variance (e.g. FP, EL, DP, RI) in abundance in the training set also displayed low sensitivity to the amount of training data. The high sensitivity of specific metabolites (e.g. lactate) and species (e.g. AC, BH) to the amount of training data indicates that further data collection would likely improve the model’s prediction performance.

**Figure 4:**
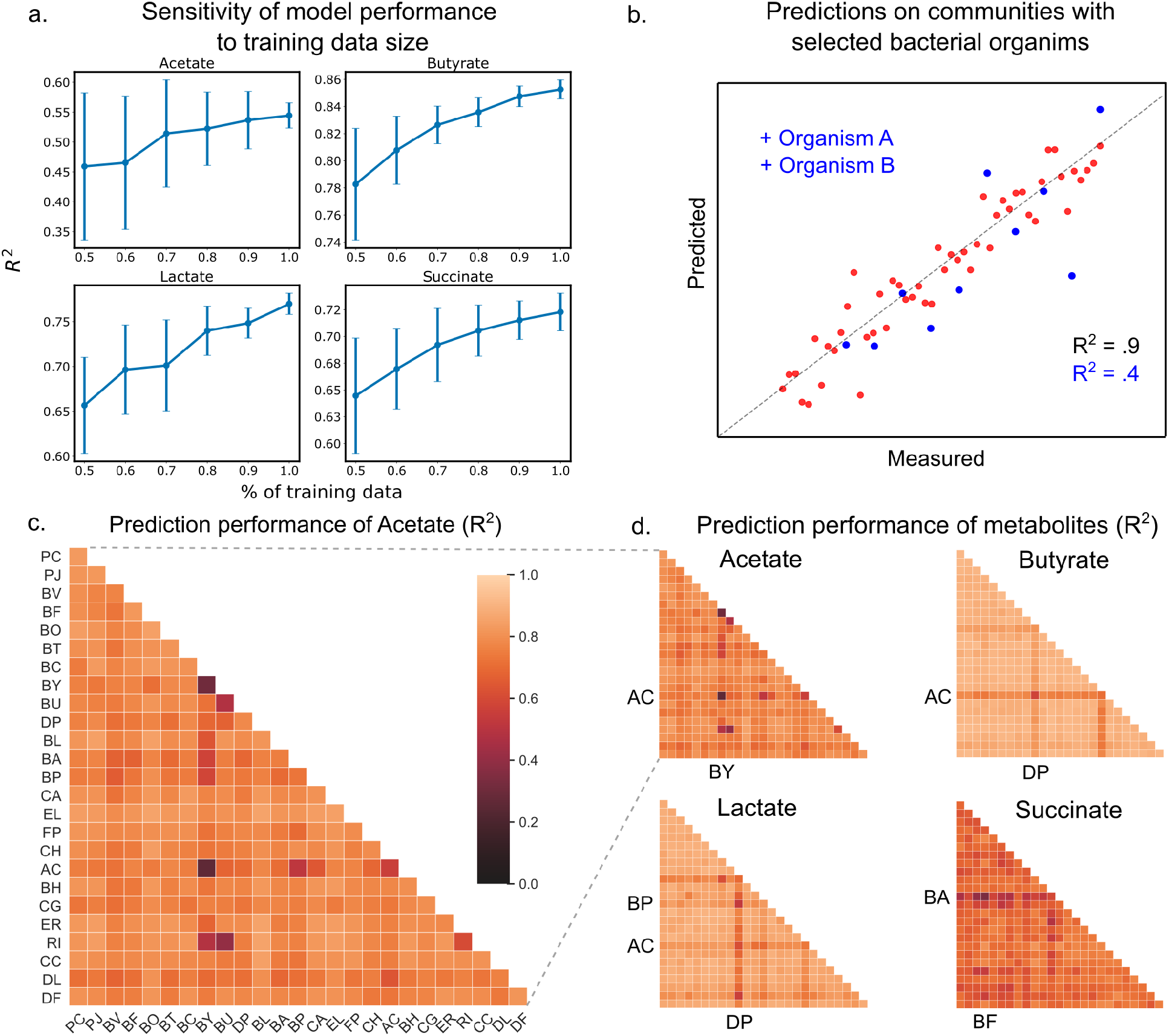
Hold-out prediction performance on sub-communities provides information about poorly understood species and interactions between species. **(a)** Sensitivity of metabolite prediction performance (*R*^2^) to the size of the training dataset. Training datasets were randomly subsampled 30 times using 50% to 100% of the total dataset in increments of 10%. Each subsampled training set was subject to 20-fold cross-validation to assess prediction performance. Lineplot of the mean prediction performance over the 30 trials for each percentage of the data. Error bars denote 1 s.d. from the mean. **(b)** Schematic scatter plot representing how communities containing species A and B define a poorly predicted subsample of the full sample set **(c)** Heatmap of prediction performance (*R*^2^) of acetate for each subset of communities containing a given species (diagonal elements) or pair of species (off-diagonal elements). **(d)** Heatmap of prediction performance for acetate, butyrate, lactate, and succinate. A sample subset containing a given species or pair of species included all communities in which the species were initially present. Predictions for each community were determined using 20-fold cross validation so that for each model the predicted samples were excluded from the training samples. N and p-values are reported in **Table S3**.

To determine how pairwise combinations of species impacted model prediction performance, we used 20-fold cross-validation to evaluate the prediction performance (*R*^2^) on subsets of the total dataset, where subsets were selected based on the presence of individual species or pairs of species (**Fig. 4b**). Using this approach, we identified individual species and species pairs that had the greatest impact on the prediction performance of metabolite concentrations. Sample subsets with poor prediction performance highlight individual species and species pairs whose presence reduces the model’s ability to make accurate predictions of final metabolite concentrations. Although the subsets were much smaller than the total data set (*n* = 761), calculation of prediction performance was not limited by small sample sizes, where the number of communities in each subset ranged from *n* = 77 to *n* = 478.

The interaction network shown in figure **Fig. 3d** shows the impact of individual species on each metabolite, but does not provide information about whether the effect is due to individual species or pairwise interactions. To determine whether pairwise interactions influence metabolite concentrations, we quantified how prediction performance changed in response to the presence individual species and pairs of species. Specifically, if prediction performance taken over a subset of communities containing a given species pair was markedly different than prediction performance for the subsets corresponding to the individual species, this suggests the pairwise interaction impacts on metabolite production. Using equation 4 (**Methods**), we found that the prediction performance of lactate and butyrate was the least sensitive to species pairs (average decrease in prediction performance for subsets with species pairs of 0.72% and 1.10% compared to corresponding single species subsets). However, the prediction performance of acetate and succinate was the most sensitive to the presence of species pairs (increase in prediction performance of 6.68% for acetate and a decrease of 2.951% for succinate). This difference in prediction performance suggests that pairwise interactions influences the production of acetate and succinate, while the production of lactate and butyrate are primarily driven by the action of individual species. The sensitivity of acetate and succinate to pairwise interactions is consistent with the inferred interaction network shown in **Fig. 3d**, which highlights multiple species-metabolite interactions for acetate and succinate and a smaller number of strong species-metabolite interactions for butyrate and lactate.

Pairs of certain *Bacteroides* and butyrate producers including BY-RI, BU-RI, and BY-AC resulted in reduced prediction performance of acetate. This suggests that interactions between specific *Bacteroides* and butyrate producers were important for acetate transformations, which is consistent with the conversion of acetate into butyrate. Based on the LIME analysis in **Fig. 3d**, AC, DP, and BP have the largest impact on lactate. Thus, the hold-out prediction performance for lactate was primarily impacted by specific pairs that include these species. In sum, these results demonstrate how the model can be used to identify informative experiments for investigating poorly understood species and interactions between species, where collection of more data would likely improve prediction performance.

### Dynamic measurements of communities reveal design rules for qualitatively distinct metabolite trajectories

We next leveraged the LSTM model’s dynamic capabilities to understand the temporal changes in metabolite concentrations and community assembly. To this end, we chose a representative subset of 95 out of the 180 communities from **Fig. 3b** (**Fig. S7a**, 60 communities for training, 34 for validation, plus the full 25 species community) and experimentally characterized species abundance and metabolite concentrations every 16 hours during community assembly (**Fig. 5a**). We analyzed the dynamic behavior of these communities using a clustering technique to extract high level design rules of species presence/absence that determined qualitatively distinct temporal metabolite trajectories (i.e. broad trends consistent across a set of communities) and exploited the LSTM framework to identify context-specific impacts of species on metabolite production (i.e. a more fine-tuned case-by-case analysis).

**Figure 5:**
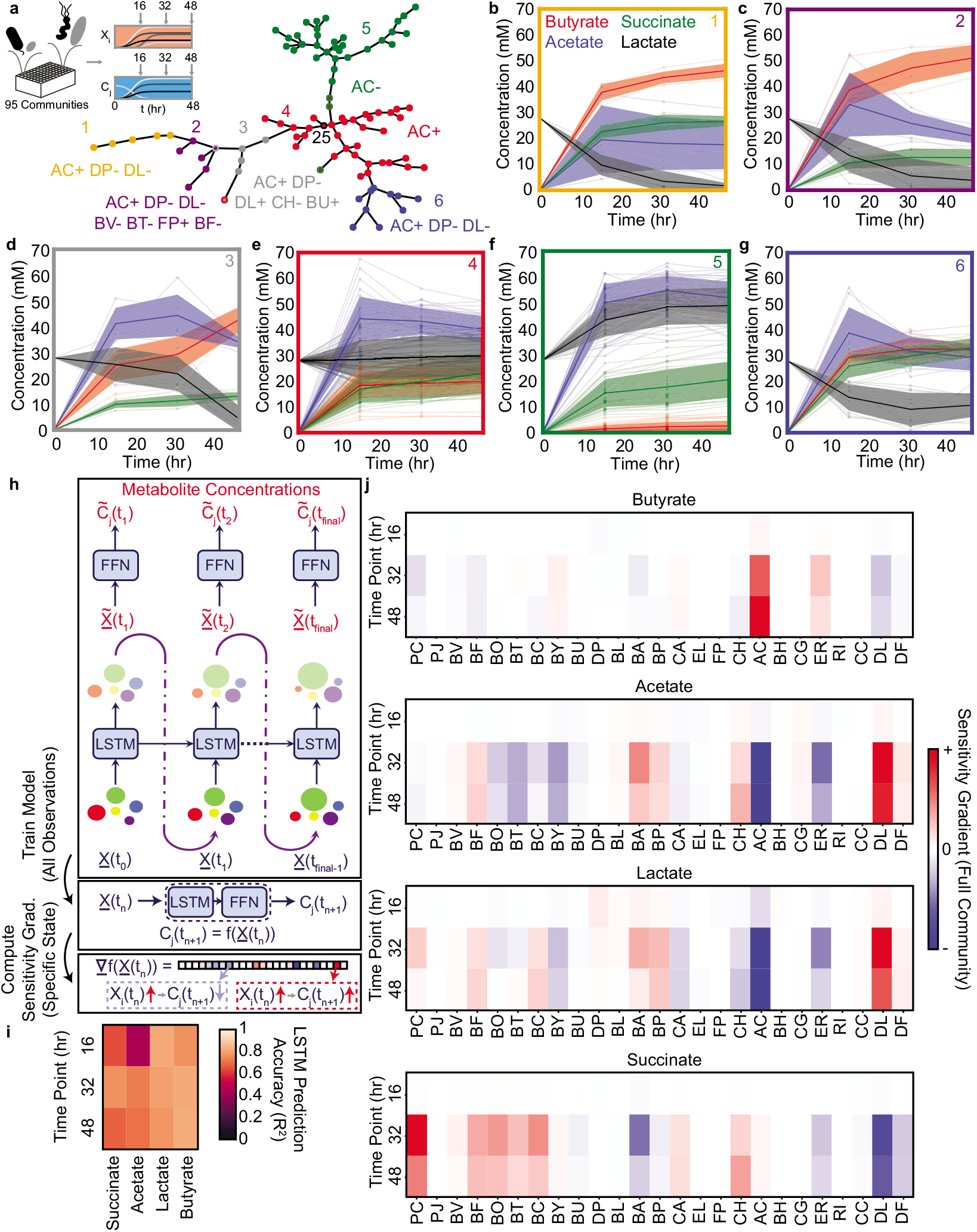
Community metabolite trajectories cluster into qualitatively distinct groups which can be classified based on presence and absence of key microbial species. **(a)** Schematic of experiment and network representing a minimal spanning tree across the 95 communities where weights (indicated by edge length) are equal to the Euclidean distance between the metabolite trajectories for each community. Node colors indicate clusters determined as described in the Methods. Red node with black outline annotated with “25” represents the community of all 25 species. Annotations indicate the most specific microbial species presence/absence rules that describe most data points in the cluster of the corresponding color as determined by a decision tree classifier (Methods). Communities that deviate from the rules for their cluster are indicated with a border matching the color of the closest cluster whose rules they do follow. Network visualization generated using the `draw_kamada_kawai` function in networkx (v2.1) for Python 3. **(b-g)** Temporal changes in metabolite concentrations for communities within each cluster (indicated by sub-plot border color), with individual communities denoted by transparent lines. Solid lines and shaded regions represent the mean plus or minus 1 s.d. of all communities in the cluster. **(h)** Schematic of LSTM model training and computation of gradients to evaluate impact of species abundance on metabolite concentrations in a specific community context. **(i)** Heatmap of model M3 prediction accuracy for four metabolites in the 34 validation communities at each time point (Pearson correlation *R*^2^. N=34 for all tests). **(j)** Heatmap of the gradient analysis of model M3 as described in (h) for the full 25-species community. N and p-values are reported in Table S3.

The temporal trajectories of species abundance and metabolite concentrations showed a wide range qualitatively distinct trends across the 95 communities (**Fig. 5b-g**). For example, some metabolites concentrations monotonically increased (e.g. butyrate in **Fig. 5b,c,e,g**), monotonically decreased (e.g. lactate in **Fig. 5b,c**) or exhibited biphasic dynamics (e.g. acetate in **Fig. 5c**). To determine if there were communities with similar temporal changes in metabolite concentrations, we clustered communities using a minimal spanning tree [32] on the Euclidean distance between the metabolite trajectories of each pair of communities (**Fig. 5a**). The resulting six clusters exhibited high quantitative within-cluster similarity and qualitatively distinct metabolite trajectories (**Fig. 5b-g**). Clusters 4 and 5 which contained the largest number of communities had a high fraction of “distributed” communities (**Fig. 3b**). Clusters with a smaller number of communities contained a higher percentage of “corner” communities (**Fig. S7b,c**). Therefore, the use of LSTM results from an initial experiment to identify “corner” communities elucidated communities with qualitatively distinct temporal behaviors. These communities were unlikely to be discovered via random sampling of sub-communities due to the high density of points towards the center of the distribution and low density in the tails of the distribution (**Fig. 3b**). Additionally, some “corner” communities that were similar in metabolite profiles when considering the end-point measurement separated into different clusters when considering the dynamic data (e.g. Clusters 2 and 3, which have similar metabolite profiles at 48 hr but qualitatively distinct dynamics (**Fig. 5b**). This demonstrates that using a community design approach to explore the extremes of system behaviors with a limited time resolution enabled the identification of additional distinct behaviors when the extreme communities were characterized with higher time resolution.

To identify general patterns in species presence/absence of these communities that could explain the temporal behaviors of each cluster, we used a decision tree analysis to identify an interpretable classification scheme (**Fig. S7d**). Using this approach, we observed that the large clusters were separated by relatively simple classification rules (i.e. AC+ for cluster 4 and AC-for cluster 5), whereas the smaller clusters had more complex classification rules involving larger combinations of species (3-7 species), all involving AC, DP, and DL (**Fig. 5a**). The influential role of DP was corroborated by a previous study showing that DP substantially inhibits butyrate production [25]. In addition, the inferred microbe-metabolite networks based on the LSTM model M2 demonstrated that the presence of DL was linked to higher acetate and lower succinate production (**Fig. 3d**), consistent with its key role in shaping metabolite dynamics in this system. The variation in the number of communities across clusters is consistent with previous observations that species-rich microbial communities tend towards similar behavior(s) (e.g. Clusters 4 and 5 contained many communities). By contrast, more complex design criteria are required to identify communities that deviate from this typical behavior (e.g. Clusters 1-3 and 6 contained few communities) [25].

While our clustering analysis identified general design rules for metabolite trajectories, there remained unexplained within-cluster variation. Thus, we used the LSTM framework to identify those effects beyond these general species presence/absence rules that determine the precise metabolite trajectory of a given community. Simultaneous predictions of species abundance and the concentration of all four metabolites at all time points necessitates specific modifications to the LSTM architecture shown in **Fig. 1a**. In particular, we consider a 29-dimensional input vector whose first 25 components correspond to the species abundance, while the remaining 4 components correspond to the concentration of metabolites (**Fig. 5h**). The 29-dimensional feature vector is suitably normalized so that the different components have zero mean and unity variance. The feature scaling is important to prevent over reliance on features with a broad range of values. The output of each LSTM unit is fed into the input block of the subsequent LSTM unit in order to advance the model forward in time. The reason behind concatenating instantaneous species abundances with metabolite concentrations can be understood as follows. Prediction of metabolite concentrations at various time points requires a time-series model (either using ODEs or LSTM in this case). Further, the future trajectory of metabolite concentrations is a function of both the species abundance, as well as the metabolite concentrations at the current time instant. Therefore, we concatenate both the metabolite concentrations and species abundances to create a 29-dimensional feature vector. The trained LSTM framework on the 60 training communities (model M3) displayed good prediction performance on the metabolite concentrations of the 34 validation communities plus the full 25-species community (**Fig. 5i**). The prediction accuracy of species abundance was lower than metabolite concentrations, presumably due to the limited number of training set observations of each species (**Fig. S8**).

We used a a gradient-based sensitivity analysis of the LSTM model M3 to provide biological insights into the contributions of each species on the temporal changes in metabolite concentrations (**Fig. 5h,j**, Methods). This method involves computing partial derivatives of output variables of interest with respect to input variables, which are readily available through a single backpropagation pass [33, 34]. As an example case, we applied this analysis approach to the full 25-species community, which was grouped into Cluster 4, with the design rule “AC+” (**Fig. 5a**). Consistent with this design rule, we observed strong sensitivity gradients between the abundance of AC and the concentrations of butyrate, acetate, and lactate, consistent with our biological understanding of the system [25]. Beyond the “AC+” design rule, there was a strong sensitivity gradient between DL and acetate and succinate, consistent with the inferred networks based on the LSTM model M2 that used single time point observations (**Fig. 3d**). Further, the contributions of certain species on metabolite production varied as a function of time. For instance, in the initial time point, species abundances were similar and thus the contribution of individual species to metabolite production is more uniform. However, interactions between species during community assembly enhanced the contribution of specific metabolite driver species such as AC. In addition, the contributions of individual species such as PC and BA to succinate production peaked at 32 hours and then decreased by 48 hours, highlighting that the effects of these species on succinate production were maximized at intermediate time points. In sum, the proposed gradient-based method identified the quantitative contributions of each species to metabolite production as a function of time for a specific case, identifying context-specific behaviors beyond the previously identified broader design rules. These two complementary approaches are useful for identifying design rules for metabolite dynamics. The clustering method can identify broad design rules for species presence/absence and the LSTM analysis approach can uncover fine-tuned quantitative contributions of species to the temporal changes in community-level functions.

## DISCUSSION

We have demonstrated that an LSTM modeling framework trained on species abundance and metabolite production in synthetic human gut communities can accurately predict multiple functions of microbial communities. This model is powerful for designing communities with target metabolite profiles. Due to its flexibility, the LSTM model outperforms the widely used gLV model in the presence of higher-order interactions. We leveraged the computational efficiency of LSTM model to predict the metabolite profiles of tens of millions of communities. We used these model predictions to identify sparsely represented “corner case” communities that maximized/minimized community-level production of four health-relevant metabolites. In the absence of a predictive model, these infrequent communities would have been difficult to discover among the vast metabolic landscape of possible communities.

Beyond the model’s predictive capabilities, we showed that biological information including significant microbe-metabolite and microbe-microbe interactions, can be extracted from LSTM models. These biological insights could enable the discovery of key species and interactions driving community functions of interest. Further, this could inform the design of microbial communities from the bottom-up or interventions to manipulate community-level behaviors. For example, the inferred microbe-metabolite network highlighted AC is a major ecological driver of several metabolites including butyrate, acetate and lactate in our system. In addition, this microbe-metabolite network did not include species of the highly abundant genus *Bacteroides* but instead featured members of Firmicutes (AC, ER, DL), Actinobacteria (BA, BP, EL), Proteobacteria DP and Bacteroidetes PC. Notably, *Bacteroides* displayed numerous interactions in the microbe-microbe interaction network, suggesting that they played a key role in the growth of constituent community members opposed to production of specific metabolites. Therefore, our model suggests that *Bacteroides* influence broad ecosystem functions such as community growth dynamics whereas species highlighted in the microbe-metabolite network contribute to specialized functions such as the production of specific metabolites [35]. Therefore, the microbe-metabolite interaction network could be used to identify key species that could be targeted for manipulating the dynamics of specific metabolites.

We performed time-resolved measurements of metabolite production and species abundance using a set of designed communities and demonstrated that communities tend towards a typical dynamic behavior (i.e. Clusters 4 and 5). Therefore, random sampling of sub-communities from the 25-member system would likely exhibit behaviors similar to Clusters 4 and 5. We used the LSTM model to identify “corner cases” communities that produce metabolite concentrations near the tails of the metabolite distributions at a single time point. Thus, the model allowed us to identify unique sub-clusters with disparate dynamic behaviors. We demonstrated that the endpoint model predictions were confirmatory (**Fig. 3c**) and also led to new discoveries when additional measurements were made in the time dimension. Specifically, we found that some “corner cases” communities identified based on prediction of a single time-point displayed distinct dynamic trajectories. For instance, Clusters 2 and 3 based on the decision tree classifier displayed similar end-point metabolite concentrations (**Fig. 5c,d**). However, lactate decreased immediately over time in Cluster 2 communities but remained high until approximately 30 hr and then decreased in Cluster 3 communities. The design rule for Cluster 3 included the presence of lactate producers BU and DL (**Fig. S4**), suggesting that these individual species’ lactate producing capabilities enabled the community to maintain a high lactate concentration for an extended period of time in the context of the Cluster 3 communities. While we focused on the production of four health-relevant metabolites produced by gut microbiota, a wide range of health-relevant compounds are produced by gut bacteria. Therefore, communities that cluster together based on dynamic trends the four measured metabolites could separate into new clusters based on the temporal patterns of other compounds produced or degraded by the communities.

Time-resolved measurements were required to reveal the different dynamic behaviors of communities in Clusters 2 and 3 to improve understanding and design of community functions. The ability to resolve differences in the dynamic trajectories of communities requires time sampling when the system behavior is changing as a function of time as opposed to time sampling once the system has reached a steady-state (i.e. saturated as a function of time). The time to reach steady-state varied across different communities and metabolites of interest. For instance, lactate reached steady-state at an earlier time point ( 12 hr) in Cluster 4 communities whereas communities in Cluster 3 approached steady-state at a later time point ( 48 hr). Therefore, model-guided experimental planning could be used to identify the optimal sampling times to resolve differences in community dynamic behaviors. The dynamic behaviors of the synthetic communities characterized *in vitro* may likely exhibit significant differences to their behaviors in new environments such as the mammalian gut. However, communities in sub-clusters whose behaviors deviated substantially from the typical community behaviors (e.g. Clusters 2 and 3 versus Clusters 4 and 5) may be more likely than random to display unique dynamic behaviors *in vivo*. Future work will investigate whether the *in vitro* dynamic behavior cluster patterns can be used as prior information to guide the design of informative communities in new environments for building predictive models.

While our current approach treated microbiome species composition as the sole set of design variables in a constant environmental background, microbiomes in reality are impacted by differences in the physicochemical composition of their environment [36]. Given sufficient observations of community behavior under varied environmental contexts (e.g. presence/absence of certain nutrients), our LSTM approach could be further leveraged to design complementary species and environmental compositions for desired microbiome functional dynamics. Further, we can leverage the wealth of biological information stored in the sequenced genomes of the constituent organisms. Integrating methods such as genome scale models [37] with our deep learning framework could leverage genomic information to enable predictions when the genomes of the organisms are varied (i.e., alternative strains of the same species with disparate metabolic capabilities). In this case, introducing variables representing the presence/absence of specific metabolic reactions would potentially enable the model to predict the impact of a species with a varied set of metabolic reactions on a given set of functions without new experimental observations. Integrating this information into the model could thus enable a mapping between genome information and community-level functions.

While previous approaches have used machine learning methods to predict microbiome functions based on microbiome species composition [18, 19], our approach is a major step forward in predicting the future trajectory of microbiome function based on an initial state of species composition. The dynamic nature of our approach enables applications to design optimal initial community compositions or interventions to perturb an existing community to achieve desired behavior in the future. The flexibility of our approach to various time resolutions is especially useful in scenarios where a microbiome may display potentially undesired transients on the path from an initial state to a desired final state. For instance, in treatment of gut microbiome dysbiosis, it is important to ensure that any transient states of the microbiome are not harmful to the host (e.g. pathogen blooms or overproduction of toxic metabolites) as the system approaches a desired healthy state [38]. However, because predictions with increased time resolution require more data for model training, the ability of our approach to work simply with initial and final observations is useful for scenarios where transient states may be less important, such as in bioprocesses where the concentration of products at the time of harvest is the key design objective [39, 40]. Finally, the computational efficiency and accuracy of the LSTM model could be exploited in the future for autonomous design and optimization of multifunctional communities via computer-controlled design-test-learn cycles [41].

## METHODS

### Strain Maintenance and Culturing

All anaerobic culturing was carried out in an anaerobic chamber with an atmosphere of 2.5 ± 0.5% H_2_, 15 ± 1% CO_2_ and balance N_2_. All prepared media and materials were placed in the chamber at least overnight before use to equilibrate with the chamber atmosphere. The strains used in this work were obtained from the sources listed in Table S1 and permanent stocks of each were stored in 25% glycerol at −80°C as previously described [25]. Batches of single-use glycerol stocks were produced for each strain by first growing a culture from the permanent stock in anaerobic basal broth (ABB) media (Oxoid) to stationary phase, mixing the culture in an equal volume of 50% glycerol, and aliquoting 400*μ*L into Matrix Tubes (ThermoFisher) for storage at −80°C. Quality control for each batch of single-use glycerol stocks included (1) plating a sample of the aliquoted mixture onto LB media (Sigma-Aldrich) for incubation at 37°C in ambient air to detect aerobic contaminants and (2) Illumina sequencing of 16S rDNA isolated from pellets of the aliquoted mixture to verify the identity of the organism. For each experiment, precultures of each species were prepared by thawing a single-use glycerol stock and combining the inoculation volume and media listed in Table S1 to a total volume of 5 mL (multiple tubes inoculated if more preculture volume needed) for stationary incubation at 37°C for the preculture incubation time listed in Table S1. All experiments were performed in a chemically defined medium (DM38), as previously described [25], the composition of which is provided in Table S2. This medium supports the individual growth of all organisms except *Faecalibacterium prausnitzii* [25].

### Community Culturing Experiments and Sample Collection

Synthetic communities were assembled using liquid handling-based automation as described previously [25]. Briefly, each species’ preculture was diluted to an OD_600_ of 0.0066 in DM38. Community combinations were arrayed in 96 deep well (96DW) plates by pipetting equal volumes of each species’ diluted preculture into the appropriate wells using a Tecan Evo Liquid Handling Robot inside an anaerobic chamber. For experiments with multiple time points, duplicate 96DW plates were prepared for each time point. Each 96DW plate was covered with a semi-permeable membrane (Diversified Biotech) and incubated at 37°C. After the specified time had passed, 96DW plates were removed from the incubator and samples were mixed by pipette. Cell density was measured by pipetting 200*μ*L of each sample into one 96 well microplate (96W MP) and diluting 20 L of each sample into 180*μ*L of PBS in another 96W MP and measuring the OD_600_ of both plates (Tecan F200 Plate Reader). We selected the value that was within the linear range of the instrument for each sample. 200*μ*L of each sample was transferred to a new 96DW plate and pelleted by centrifugation at 2400xg for 10 minutes. A supernatant volume of 180*μ*L was removed from each sample and transferred to a 96-well microplate for storage at −20°C and subsequent metabolite quantification by high performance liquid chromatography (HPLC). Cell pellets were stored at −80°C for subsequent genomic DNA extraction and 16S rDNA library preparation for Illumina sequencing. 20*μ*L of each supernatant was used to quantify pH using a phenol Red assay [42]. Phenol red solution was diluted to 0.05% weight per volume in 0.9% w/v NaCl. Bacterial supernatant (20*μ*L) was added to 180*μ*L of phenol red solution in a 96W MP, and absorbance was measured at 560 nm (Tecan Spark Plate Reader). A standard curve was produced by fitting the Henderson-Hasselbach equation to fresh media with a pH ranging between 3 to 11 measured using a standard electro-chemical pH probe (Mettler-Toledo). We used (1) to map the pH values to the absorbance measurements.

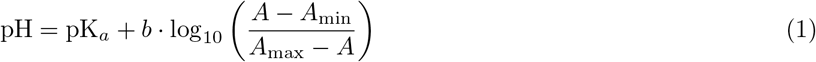

The parameters *b* and pK_*a*_ were determined using a linear regression between pH and the log term for the standards in the linear range of absorbance (pH between 5.2 and 11) with *A*_max_ representing the absorbance of the pH 11 standard, *A*_min_ denoting the absorbance of the pH 3 standard and *A* representing the absorbance of each condition.

### HPLC Quantification of Organic Acids

Butyrate, succinate, lactate, and acetate concentrations in culture supernatants were quantified as described previously [25]. Supernatant samples were thawed in a room temperature water bath before addition of 2*μ*L of H_2_SO_4_ to precipitate any components that might be incompatible with the running buffer. The samples were then centrifuged at 2400xg for 10 minutes and then 150*μ*L of each sample was filtered through a 0.2*μ*m filter using a vacuum manifold before transferring 70*μ*L of each sample to an HPLC vial. HPLC analysis was performed using a Shimadzu HPLC system equipped with a SPD-20AV UV detector (210 nm). Compounds were separated on a 250 × 4.6 mm Rezex^©^ ROA-Organic acid LC column (Phenomenex Torrance, CA) run with a flow rate of 0.2 ml min^−1^ and at a column temperature of −50°C. The samples were held at 4°C prior to injection. Separation was isocratic with a mobile phase of HPLC grade water acidified with 0.015 N H_2_*SO*_4_ (415*μ*LL^−1^). At least two standard sets were run along with each sample set. Standards were 100, 20, and 4 mM concentrations of butyrate, succinate, lactate, and acetate, respectively. The injection volume for both sample and standard was 25*μ*l. The resultant data was analyzed using the Shimadzu LabSolutions software package.

### Genomic DNA Extraction and Sequencing Library Preparation

Genomic DNA extraction and sequencing library preparation were performed as described previously [25]. Genomic DNA was extracted from cell pellets using a modified version of the Qiagen DNeasy Blood and Tissue Kit protocol. First, pellets in 96DW plates were removed from −80°C and thawed in a room temperature water bath. Each pellet was resuspended in 180*μ*L of enzymatic lysis buffer (20 mM Tris-HCl (Invitrogen), 2 mM Sodium EDTA (Sigma-Aldrich), 1.2% Triton X-100 (Sigma-Aldrich), 20 mg/mL Lysozyme from chicken egg white (Sigma-Aldrich)). Plates were then covered with a foil seal and incubated at 37°C for 30 minutes with orbital shaking at 600 RPM. Then, 25*μ*L of 20mgmL^−1^ Proteinase K (VWR) and 200 L of Buffer AL (QIAGEN) were added to each sample before mixing with a pipette. Plates were then covered by a foil seal and incubated at 56°C for 30 minutes with orbital shaking at 600 RPM. Next, 200*μ*L of 100% ethanol (Koptec) was added to each sample before mixing and samples were transferred to a Nucleic Acid Binding (NAB) plate (Pall) on a vacuum manifold with a 96DW collection plate. Each well in the NAB plate was then washed once with 500*μ*L Buffer AW1 (QIAGEN) and once with 500*μ*L of Buffer AW2 (QIAGEN). A vacuum was applied to the Pall NAB plate for an additional 10 minutes to remove any excess ethanol. Samples were then eluted into a clean 96DW plate from each well using 110*μ*L of Buffer AE (QIAGEN) preheated to 56°C. Genomic DNA samples were stored at −20°C until further processing.

Genomic DNA concentrations were measured using a SYBR Green fluorescence assay and then normalized to a concentration of 1ngL^−1^ by diluting in molecular grade water using a Tecan Evo Liquid Handling Robot. First, genomic DNA samples were removed from −20°C and thawed in a room temperature water bath. Then, 1*μ*L of each sample was combined with 95*μ*L of SYBR Green (Invitrogen) diluted by a factor of 100 in TE Buffer (Integrated DNA Technologies) in a black 384-well microplate. This process was repeated with two replicates of each DNA standard with concentrations of 0, 0.5, 1, 2, 4, and 6ngL^−1^. Each sample was then measured for fluorescence with an excitation/emission of 485/535 nm using a Tecan Spark plate reader. Concentrations of each sample were calculated using the standard curve and a custom Python script was used to compute the dilution factors and write a worklist for the Tecan Evo Liquid Handling Robot to normalize each sample to 1ngL^−1^ in molecular grade water. Samples with DNA concentration less than 1ngL^−1^ were not diluted. Diluted genomic DNA samples were stored at −20°C until further processing.

Amplicon libraries were generated from diluted genomic DNA samples by PCR amplification of the V3-V4 of the 16S rRNA gene using custom dual-indexed primers for multiplexed next generation amplicon sequencing on Illumina platforms [25, 43]. Primers were arrayed in skirted 96 well PCR plates (VWR) using an acoustic liquid handling robot (Labcyte Echo 550) such that each well received a different combination of one forward and one reverse primer (0.1*μ*L of each). After liquid evaporated, dry primers were stored at −20°C. Primers were resuspended in 15*μ*L PCR master mix (0.2*μ*L Phusion High Fidelity DNA Polymerase (Thermo Scientific), 0.4*μ*L 10 mM dNTP Solution (New England Biolabs), 4*μ*L 5x Phusion HF Buffer (Thermo Scientific), 4*μ*L 5M Betaine (Sigma-Aldrich), 6.4*μ*L Water) and 5*μ*L of normalized genomic DNA to give a final concentration of 0.05 M of each primer. Primer plates were sealed with Microplate B seals (Bio-Rad) and PCR was performed using a Bio-Rad C1000 Thermal Cycler with the following program: initial denaturation at 98°C (30 s); 25 cycles of denaturation at 98°C (10 s), annealing at 60°C (30 s), extension at 72°C (60 s); and final extension at 72°C (10 minutes). 2*μ*L of PCR products from each well were pooled and purified using the DNA Clean & Concentrator (Zymo) and eluted in water. The resulting libraries were sequenced on an Illumina MiSeq using a MiSeq Reagent Kit v3 (600-cycle) to generate 2×300 paired end reads.

### Bioinformatic Analysis for Quantification of Species Abundance

Sequencing data were used to quantify species relative abundance as described previously [25, 43]. Sequencing data were demultiplexed using Basespace Sequencing Hub’s FastQ Generation program. Custom python scripts were used for further data processing as described previously [25, 43]. Paired end reads were merged using PEAR (v0.9.10) [44] after which reads without forward and reverse annealing regions were filtered out. A reference database of the V3-V5 16S rRNA gene sequences was created using consensus sequences from next-generation sequencing data or Sanger sequencing data of monospecies cultures. Sequences were mapped to the reference database using the mothur (v1.40.5) [45] command classify.seqs (Wang method with a bootstrap cutoff value of 60). Relative abundance was calculated as the read count mapped to each species divided by the total number of reads for each condition. Absolute abundance of each species was calculated by multiplying the relative abundance by the OD_6_00 measurement for each sample. Samples were excluded from further analysis if .1% of the reads were assigned to a species not expected to be in the community (indicating contamination).

### Long Short Term Memory for dynamic prediction on Microbial Communities

Long short term memory (LSTM) networks belong to the class of recurrent neural networks (RNNs) and model time-series data. They were first introduced by Hochreiter et al. [46] to overcome the vanishing or exploding gradients problem [47] that occur due to long-term temporal dependencies. Since their inception, LSTMs have been further refined [48, 49] and find numerous applications in several domains, including but not limited to neuroscience [50], weather forecasting [51], predictive finance [52], Google Voice for speech recognition [53, 52] and Google Allo for message suggestion [54].

Similar to any recurrent neural network, an LSTM network, too, comprises of a network of multiple LSTM units, each representing the input-output map at a time instant. Fig. 1 shows the schematic of the proposed LSTM network architecture for abundance prediction. For a microbial community comprising of *N* species, each LSTM unit models the dynamics at time *t* using the following set of equations:

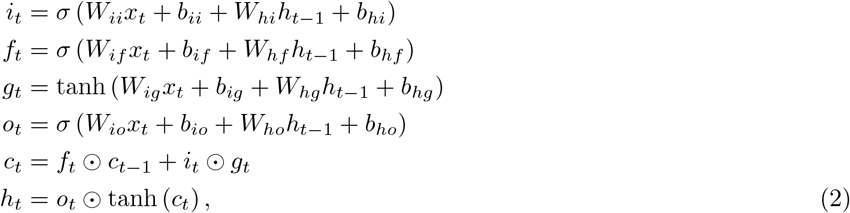

where *h_t_, c_t_, x_t_* are the hidden state, cell state and input abundance at time *t*, respectively, and *i_t_, f_t_, g_t_, o_t_* are input, forget, cell and output gates, respectively. *σ* is the sigmoid function, and ⊙ denotes the Hadamard product. The parameters {*W_mn_, b_mn_*} for *m, n* ∈ {*f, g, h, i, o*} are trainable and shared across all LSTM units. The output gate *o_t_* is further used to generate the abundance for next time instant as:

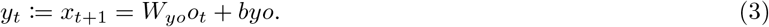

As shown in Fig. 1, *y_t_* is fed to the LSTM unit at the next timestep (*t* + 1), which in turn predicts the species abundance at time *t* + 2. The process is repeated across multiple LSTM units in order to obtain *x*_*t*_final__. The entire architecture is trained to minimize the mean-squared loss between the predicted abundance *x*_*t*_final__ and true abundance 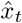.

### Using Teacher forcing for intermittent time-series forecasting

The end-goal for the proposed LSTM-network based abundance predictor is to accurately capture the steady-state (final) abundance from initial abundance. In typical LSTM networks, the output of the recurrent unit at the previous timestep *y*_*t*−1_ is used as an input to the recurrent unit at the current timestep *x_t_*. This kind of recurrent model, while has the ability to predict final abundance, is incapable to handle he one-step-ahead prediction. The problem is even more critical when one tries to anticipate more than a single timestep into the future. *Teacher forcing* [55] entails a training procedure for recurrent networks, such as LSTMs, where ‘true’ abundances at intermittent timesteps are used to guide (like a teacher) the model to accurately anticipate one-step-ahead abundance.

Teacher forcing is an efficient method of training RNN models that use the ground truth from a prior time step as input. This is achieved by occasionally replacing the predicted abundance *y*_*t*−1_ from the previous timestep with the true abundance 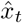 at the current timestep as input abundance to the LSTM unit at the current timestep during the training process. Teacher forcing not only stabilizes the training process, it *forces* the output abundances at all times to closely match the corresponding true abundances. This is precisely why we do not just use the ground truth abundances at intermittent timesteps in order to robustify the prediction of steady-state abundance. Once trained, the inference in such models is achieved by ignoring the ground truth abundances and using the predicted abundance from previous instant to roll forward the model in time.

### Metabolite Profiling

Microbial communities are a rich source of a variety of metabolites that are very commonly used as nutritional supplements, natural compounds to cure infectious diseases and in sustainable agriculture development. The concentration and chemical diversities of metabolites produced in a microbial community is a direct consequence of the diversity of interactions between organisms in the community. In essence, the dynamical evolution of relative species abundance and intra-community interactions govern the nature and amount of metabolites produced in the community. The functional map between species abundance and concentration of metabolites is highly complex and nonlinear, and is often approximated using simple regressors involving unary and pairwise interaction terms. In this paper, we model the species-metabolite map through appropriate modification of the LSTM network.

The aforementioned LSTM network for predicting the species abundance is suitably modified to augment four additional components that correspond to the concentration of metabolites at each time instant. In particular, the species abundance data (of size *N*_species_) is concatenated with the metabolite concentration data (of size *N*_metabs_) to form a (*N*_species_ + *N*_metabs_)-dimensional feature vector, which is suitably normalized so that the different components have zero mean and unity variance. The feature scaling is important to prevent over reliance on features with a broad range of values. Concatenation of species abundance data and the metabolite concentration data ensures that the future trajectory of metabolite concentrations evolves as a function of both the species abundance, as well as the metabolite concentrations at previous time instants. As before, the (*N*_species_ + *N*_metabs_)-dimensional output of each LSTM unit is fed into the input block of the subsequent LSTM unit in order to advance the model forward in time. The model predictions at each time point is then transformed back to the original scale in order to obtain the Pearson *R*^2^ scores on the unnormalized data. Compared with existing approaches that employ ordinary differential equations (ODEs) and multiple linear regression models for predicting metabolites, the proposed architecture enables more accurate and rapid estimation of all four metabolites. All the LSTM models were implemented in Python using PyTorch on an Intel i7-7700HQ CPU @2.80GHz processor with 16GB RAM and NVIDIA GeForce GTX 1060 (6GB GDDR5) GPU. The exact details of the neural network architecture consisting of number of layers, learning rate, choices of optimizer and nonlinear activations are described in Table S2.

### Using LSTM Model to Design Multifunctional Communities

We used the LSTM model trained on previous data (Fig. 3a) to design two sets of communities: a “distributed” community set and a “corner” community set. For the “distributed” community set, we first took the predicted metabolite concentrations for all communities with .10 species and used *k*-means clustering with *k* = 100 (Python 3, scikit-learn v0.23.1, `sklearn.cluster.Kmeans` function) to identify 100 cluster centroids that were distributed across all of the predictions. We then found the closest community to each centroid in terms of Euclidean distance in the 4-dimensional metabolite concentration space. These 100 communities constituted the “distributed” community set.

For the “corner” community set, we first defined 4 “corners” in the lactate and butyrate concentration space by binning all communities with .10 species as shown in Fig. 3b:

1. 5% lowest lactate concentration communities, then 5% lowest butyrate concentration of those
2. 5% lowest lactate concentration communities, then 5% highest butyrate concentration of those
3. 5% lowest butyrate concentration communities, then 5% lowest lactate concentration of those
4. 5% lowest butyrate concentration communities, then 5% highest lactate concentration of those

Within each of those four “corners”, we identified four “sub-corners” in the acetate and succinate concentration space by binning communities as shown in Fig. 3b:

1. 5% lowest acetate concentration communities, then 5% lowest succinate concentration of those
2. 5% lowest acetate concentration communities, then 5% highest succinate concentration of those
3. 5% lowest succinate concentration communities, then 5% lowest acetate concentration of those
4. 5% lowest succinate concentration communities, then 5% highest acetate concentration of those

This process resulted in 16 “sub-corners” total. For each “sub-corner”, we then chose a random community and then identified 4 more communities that were maximally different from that community in terms of which species were present (Hamming distance). This overall process resulted in 80 communities constituting the “corner” community set.

### Composite Model: gLV Model for Predicting Species Abundance

To benchmark the performance of the LSTM model for predicting metabolite production, we used a previously described Composite Model consisting of a generalized Lotka-Volterra (gLV) model for predicting species abundance dynamics and a regression model with interaction terms to predict metabolite concentration at a given time from the species abundances at that time [25]. Because our LSTM model was trained on the same dataset as Composite Model M3 from [25], we used those gLV model parameters.

### Composite Model: Regression Models for Predicting Metabolite Concentrations

We used a Bayesian regression model for identifying the gLV parameters with interaction terms based on [56]. Our implementation is similar to the model described in [25] for predicting metabolite concentration from community composition at a particular time. However, because the regression model from [25] was focused specifically on the prediction of butyrate, we adapted the approach to prediction of multiple metabolites. First, we modified the model form to include first order and interaction terms for all 25 species, rather than just the butyrate producers. Then, we trained 4 regression models, one for each metabolite (butyrate, lactate, acetate, succinate), using the measured species abundance and measured metabolite concentrations from the same dataset used to train the LSTM model. We trained these models as described previously [25] by using Python scikit-learn [57] to perform L1 regularization to minimize the number of nonzero parameters. Regularization coefficients were chosen by using 10-fold cross validation and choosing the coefficient value with the lowest median mean-squared error across the training splits.

### Composite Model: Simulations for Prediction

Custom MATLAB scripts were used to predict community assembly using the gLV model as described previously [25]. For each community, the growth dynamics were simulated using each parameter set from the posterior distribution of the gLV model parameters. The resulting community compositions for each simulation at 48 hours were used as an input to the Python regression models to predict the concentration of each metabolite in each community for each gLV parameter set. Because of the large number of communities and the large number of parameter sets (i.e., hundreds of simulations per community), we used parallel computing (MATLAB parfor) to complete the simulations in a reasonable timeframe (~1 hr for the communities in Figure S3a).

### Understanding Relationships Between Variables

#### Using LIME

Black-box methods, such as the LSTM-networks employed in this manuscript, do not offer much insights into the underlying mechanics that make them so powerful. Consequently, any potential pitfalls that may come along with building such models remain unexplored. For networks that are of significant biological importance, basing assumptions on falsehoods can be catastrophic. We overcome this limitation by resorting to Local Interpretable Model-Agnostic Explanations (LIME) [26].

LIME has three key components: (a) *Local*, i.e., any explanation reflects the behavior of a classifier around the sampled instance, (b) *Interpretability*, i.e., the explanations offered by LIME are interpretable by human, (c) *Model-Agnostic*, i.e., LIME does not require to peak into any model. It generates explanations by analyzing the model’s behavior for an input perturbed around its neighborhood. In this manuscript, we employ LIME to explain both qualitatively and quantitatively, as to how the abundances of various species affect the concentrations of all four metabolites, and if the presence or absence of a given species has any significance on the resulting metabolite profile.

We carried out the LIME analysis to generate interpretable prediction explanations for model M2 for each community instance used to train the model. We used lime v0.2.0.1 for Python 3 (https://github.com/marcotcr/lime) to train an explainer on the predictions of the training instances for each output variable (25 species, 4 metabolites) and then generated explanation tables for every input variable (species presence/absence) for every training instance. We then determined the median value for which the presence of a given species explained the prediction for each output variable to generate the networks in Fig. 3d,e.

#### Using Prediction Sensitivity

For each metabolite (Acetate, Butyrate, Lactate, Succinate), fractions of .5, .6, .7, .8, .9, and 1 of the total dataset were randomly sampled. Each sub-sampled dataset was subject to 20-fold cross validation to determine the sensitivity of held-out prediction performance to the amount of data available for training. This process was repeated 30 times, and the average prediction over the 30 trials was used to compute the final held-out prediction performance (*R*^2^).

The sensitivity of the model to the presence of individual species and pairs of species was determined by evaluating prediction performance (*R*^2^) for subsets of the data containing each species and each possible pair of species. To evaluate how prediction performance of each metabolite was affected by the presence of species pairs, we computed the average percent difference between prediction performance taken over subsets containing a single species and all pairs of species using the following equation,

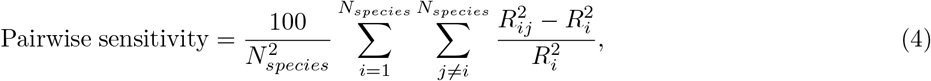

where 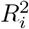 is the prediction performance taken over the subset of samples containing species *i*, and 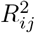 is the prediction performance taken over the subset of samples containing species *i* and *j*.

#### Using Sensitivity Gradients

Interpretability of neural-network (NN) models continues to be an interesting challenge in machine learning. While LIME is a great tool to explain what machine learning classifiers are doing, it is model-agnostic and uses simple linear models to approximate local behavior. Model-agnostic characteristic enforces retraining linear models on the training data and analyzing local perturbations, before LIME can be used to invoke interpretability. Moreover, the type of modifications that need to be performed on the data to get proper explanations are typically use case specific. Consequently, model-aware interpretability methods that take into account the weights of an already trained NN are more suitable.

For tasks, such as classification of images and videos, there is a natural way to interpret NN models using class activation maps (CAMs) [58]. CAMs assigns appropriate weighting to different convolutional filters and highlights part of the images that activate a given output class the most. However, CAMs do not extend to other NN architectures, such as LSTMs. Fortunately for us, the answer to interpretability lies in the model training itself. Let *Y* be the output variable of interest whose perturbation with respect to an input *x* needs to be estimated. The effect of *x* on *Y* can be approximated through the partial derivative 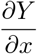. For instance, *Y* may denote butyrate concentration in an experiment, while *x* can be used to represent abundance of one of the species. The sign of the partial derivative depicts positive (or negative) correlation between the two variables, while the magnitude represents the extent of it. In order to evaluate the partial derivatives, we freeze the weights of the already trained LSTM model and declare the inputs to be variables. A single backpropagation pass then evaluates the partial derivatives of an output variable of interest with respect to all the input variables.

### Clustering Metabolite Trajectories

To generate the clusters from the dynamic community observations (Fig. 5), we used a graph-theoretic divisive clustering algorithm [59] based on the minimal spanning tree [60]. We first generated an undirected graph wherein each node was a community observed in our experiment and each edge weight was the Euclidean distance between two communities based on all metabolite measurements (4 metabolites × 3 time points=12-dimensional space for Euclidean distance calculation). We then determined the minimal spanning tree for this graph using the `minimum_spanning_tree` function in networkx (v2.1) for Python 3. We then used this minimal spanning tree to generate clusters by iteratively removing the edge with the largest weight until 6 clusters were formed. In each iteration, if any edge removal resulted in a cluster with <5 communities (i.e. minimum cluster size), that edge was returned and the next largest edge was removed. The number of clusters and minimum cluster size were chosen based on an elbow method [61], wherein scatter plots were made of the mean intracluster distance versus the number of clusters for various minimum cluster sizes and a combination of minimum cluster size and number of clusters that fell on the elbow of the plot was chosen.

### Decision Tree Classification of Metabolite Trajectories

The decision tree shown in Fig. S5d and used to produce the annotations in Fig. 5a was generated using the DecisionTreeClassifier with the default parameter settings in scikit-learn (v0.23.1) for Python 3 (visualization generated using `plot_tree` function from the same).

### Choice of Sample Sizes

Sample sizes were chosen based on limitations of experimental throughput as increased number of biological replicates would have reduced the number of possible different communities that could be observed. We chose a minimum of 2 biological replicates (for complex communities in our validation set) and some sample types have up to 7 biological replicates (such as the full community, which was repeated in most experiments as a control for consistency between experimental days).

## DATA & CODE AVAILABILITY

Pytorch implementation of the proposed LSTM model and the accompanying measurements of community composition and metabolite concentrations will be available from GitHub. The raw Illumina sequencing data will be available from Zenodo. These datasets and codes will be available at the time of submission of the revised manuscript.

## ACKNOWLEDGEMENTS

This research was supported by funding from the Army Research Office (ARO) grant number W911NF1910269 and the National Institutes of Health under grant number R35GM124774. RLC was supported in part by an NHGRI training grant to the Genomic Sciences Training Program (T32 HG002760).

## AUTHOR CONTRIBUTIONS

O.S.V., A.O.H., R.L.C. and M.B. conceived the study. R.L.C. carried out experiments. R.L.C., M.B., J.T. and Z.S. performed computational work. O.S.V., A.O.H., R.L.C., M.B. and J.T. wrote and revised the paper. O.S.V. and A.O.H. secured funding.

## Supplemental Figures

**Figure S1.**
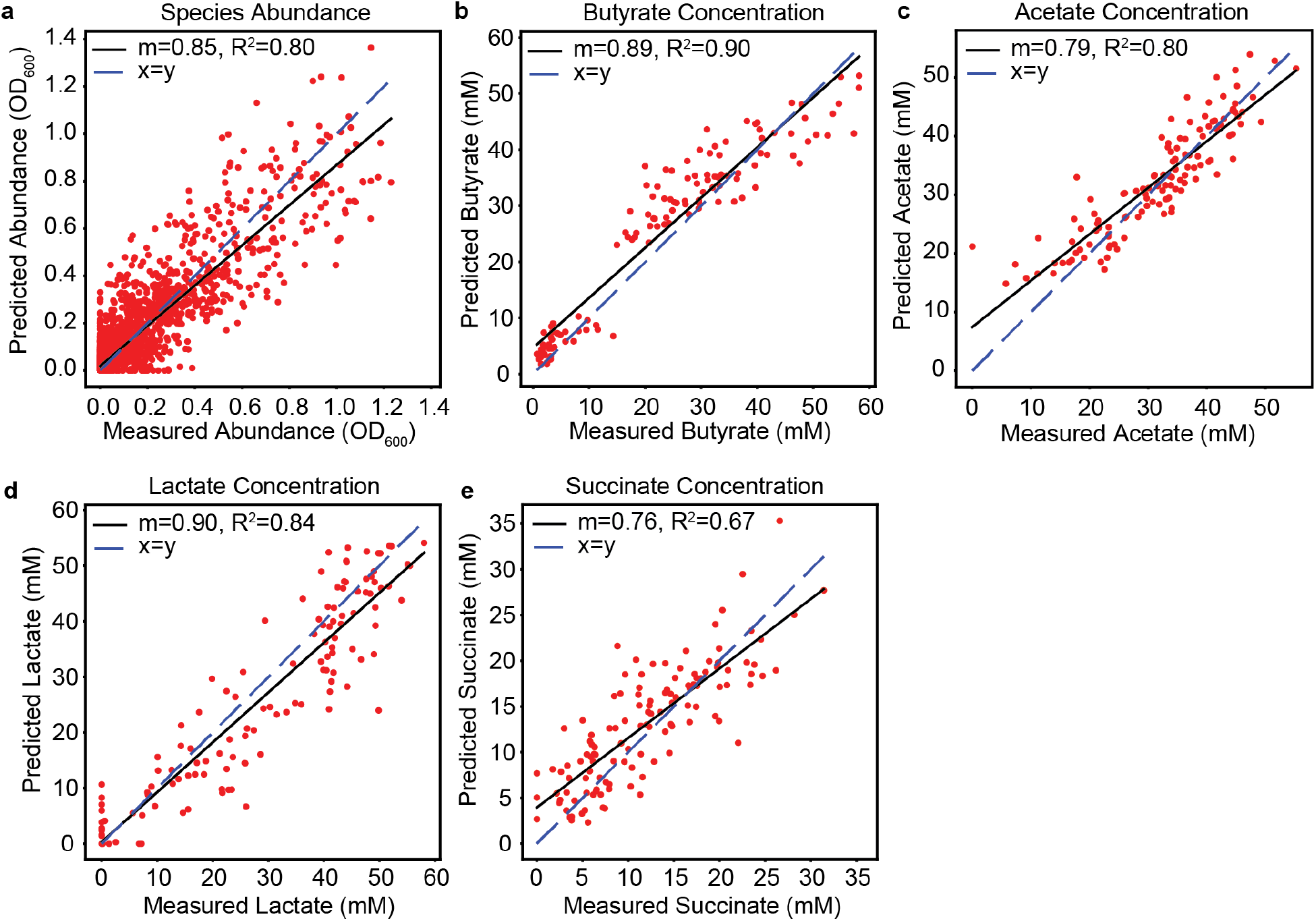
Cross-validation of LSTM model M1 predictions of species abundance and metabolite concentration. Each plot indicates the comparison of predicted versus measured species abundance (N = 1736, p = 2.64e-273) (**a**), butyrate concentration (N = 124, p = 6.19e-13) (**b**), acetate concentration (N = 124, p = 2.58e-30) (**c**), lactate concentration (N = 124, 2.67e-28) (**d**), or succinate concentration (N = 124, p = 3.34e-9) (**e**) for cross-validation of model M1 predictions of the validation communities from Clark et al., *Nature Communications*, 2021 (model trained on 110 pairwise communities, 156 communities with 3-5 species, and 124 communities with 11-17 species; cross-validation shown is prediction of a different set of 124 communities with 11-17 species, including 82 communities with all 5 butyrate producers *A. caccae, E. rectale, F. prausnitzii, C. comes and R. intestinalis* and 42 communities with the 4 butyrate producers other than *A. caccae*). Each data point indicates the average of biological replicates of a single community. Black lines indicate linear regressions with slope (m) and R^2^ indicated in the legends. Dashed blue line indicates x=y.

**Figure S2.**
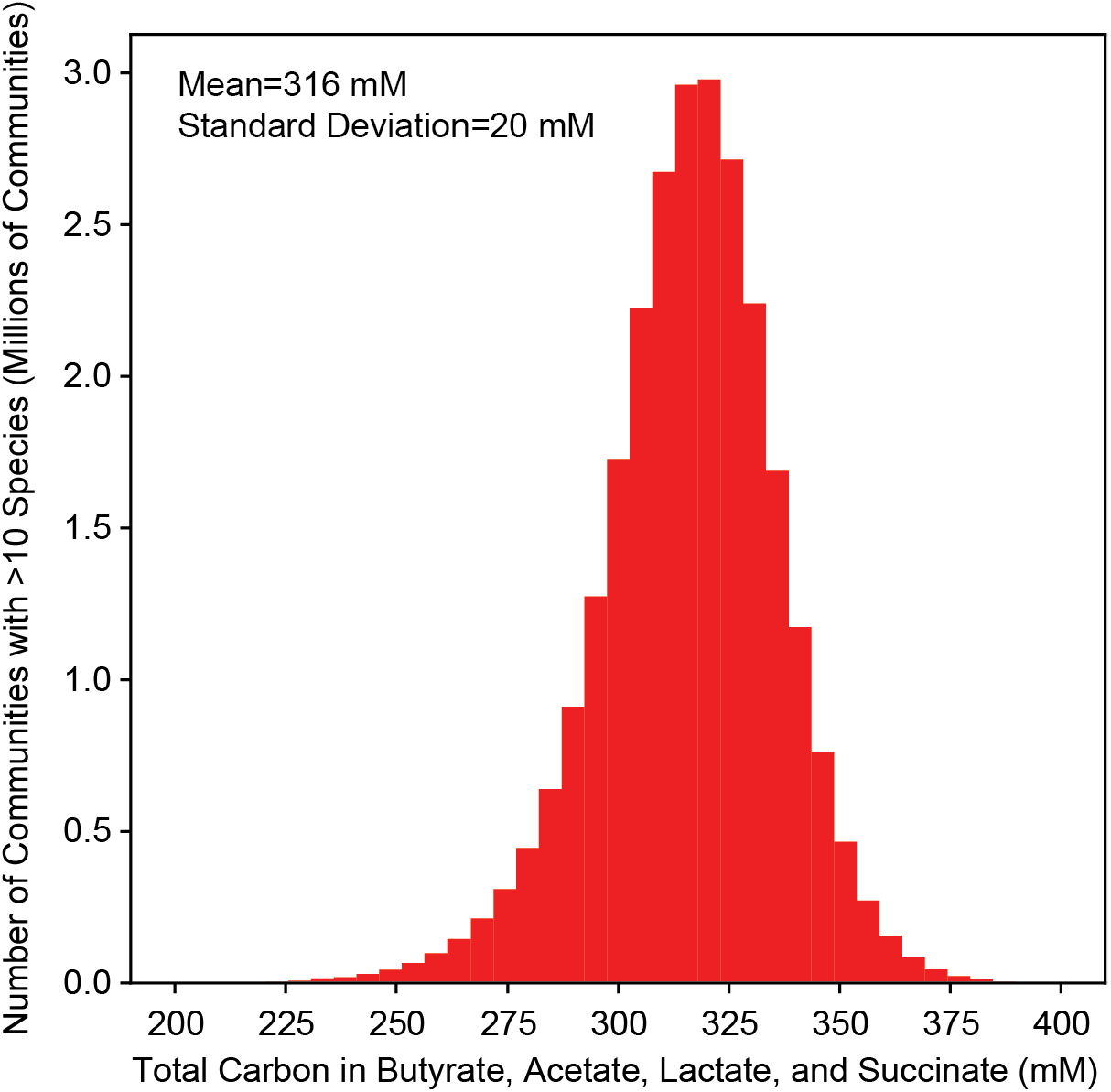
Predicted total carbon in fermentation products. Histogram of the model M1 predicted total carbon concentration in butyrate, acetate, lactate, and succinate for all possible communities with >10 species (26,434,916 communities).

**Figure S3.**
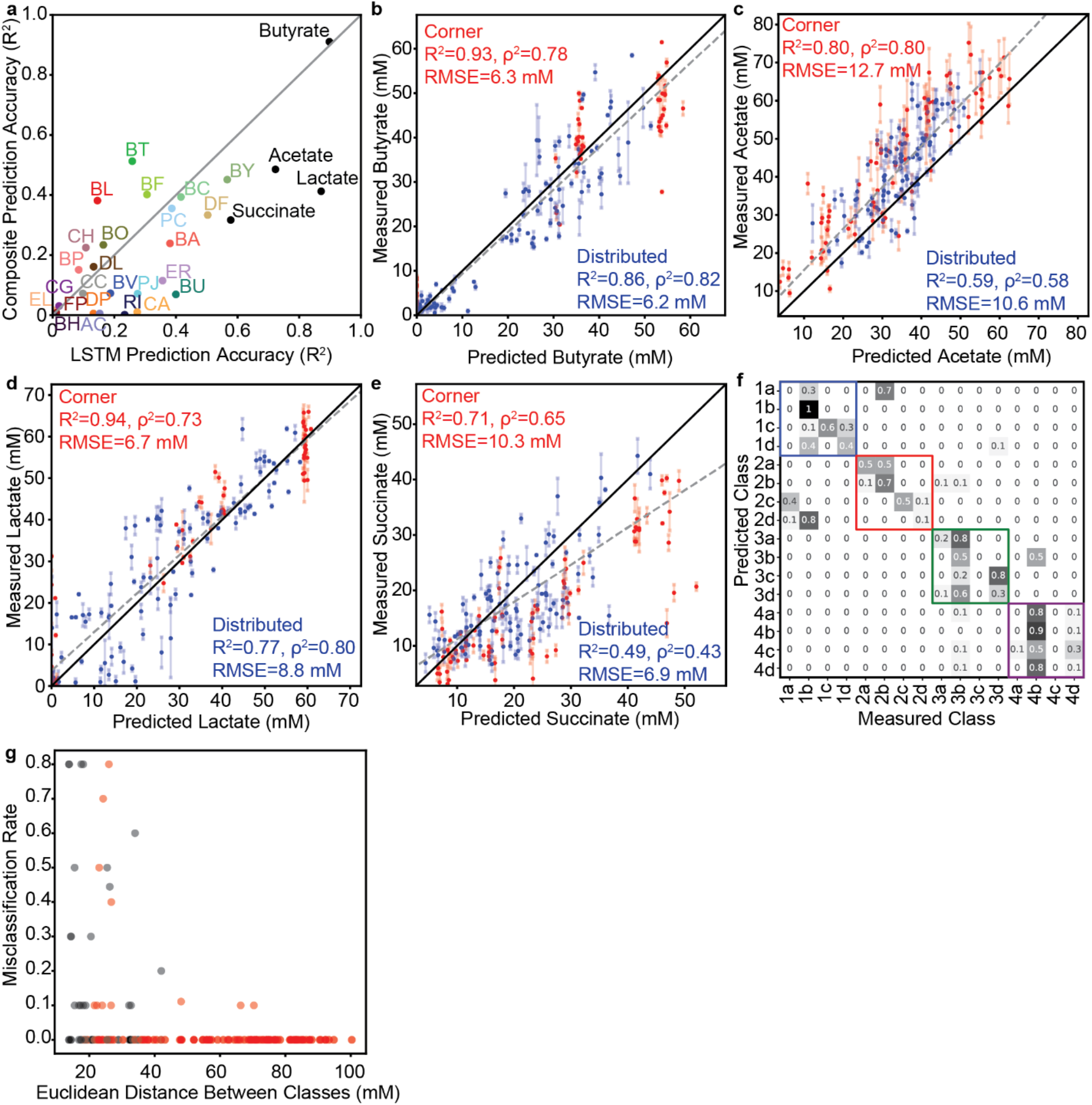
Prediction and classification statistics for model M1 predictions of designed community sets. (**a**) Scatter plot of R^2^ for prediction accuracy (correlation of predicted versus measured) of each variable (25 species abundances, 4 metabolite concentrations) by LSTM model M1 versus the Composite Model based on the method from Clark et al., *Nature Communications*, 2021. N and p-values are reported in **Table S3**. (**b-e**) Prediction accuracy of model M1 for the indicated metabolites. Dashed line indicates the linear regression for all data points. Legends indicates the R^2^, Spearman ρ^2^, and RMSE for communities from the “corner” set (red) or “distributed” set (blue) for each variable. Solid black lines indicate x=y. (**f**) Confusion matrix for classification of the “corner” communities into their specified classes (shown in **Figure 3b**). Values indicate the fraction of communities from each predicted class whose metabolite concentrations were closest (Euclidean distance) to the centroid of each class (Measured Class). Colored boxes indicate “sub-classes” that fall within the 4 major classes determined in the lactate and butyrate concentration space as shown in **Figure 3b**. (**g**) Scatter plot of misclassification rate between each pair of classes (values from **f,** fraction of communities misclassified from one class to the other) versus the Euclidean distance between the centroids of that pair of classes.

**Figure S4.**
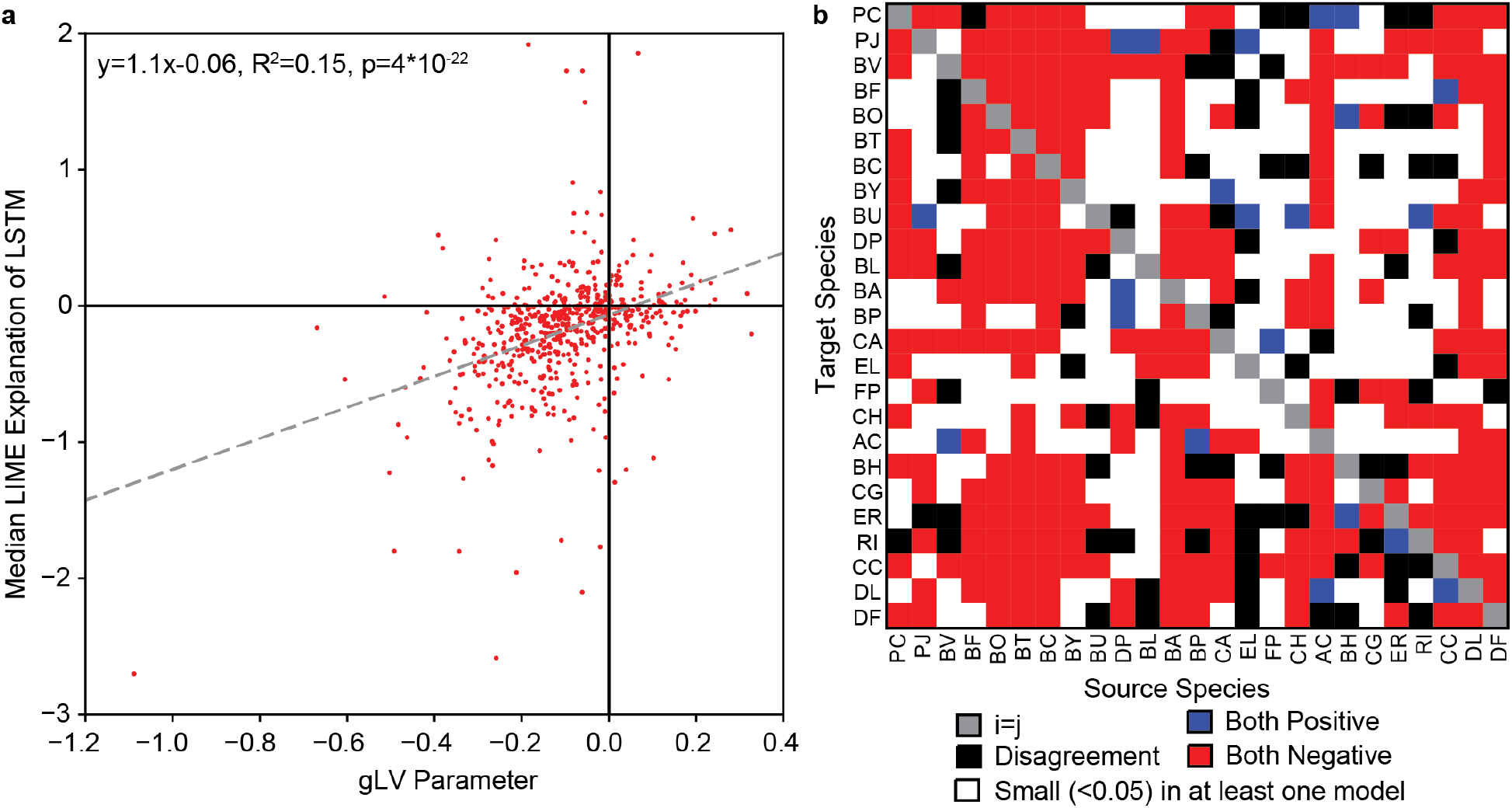
Comparison of LIME explanations of LSTM to gLV Parameters. **(a)** Scatter plot of LIME explanations of each species impact on each other species in model M2 versus the corresponding interspecies interaction parameter (a_ij_) from the gLV model from Clark et al., *Nature Communications*, 2021. Dashed line indicates the linear regression with the regression parameters shown in the legend. **(b)** Heatmap representation of qualitative agreement/disagreement between specific inter-species interactions for the same comparison as in (a). Legend describes what each color represents.

**Figure S5.**
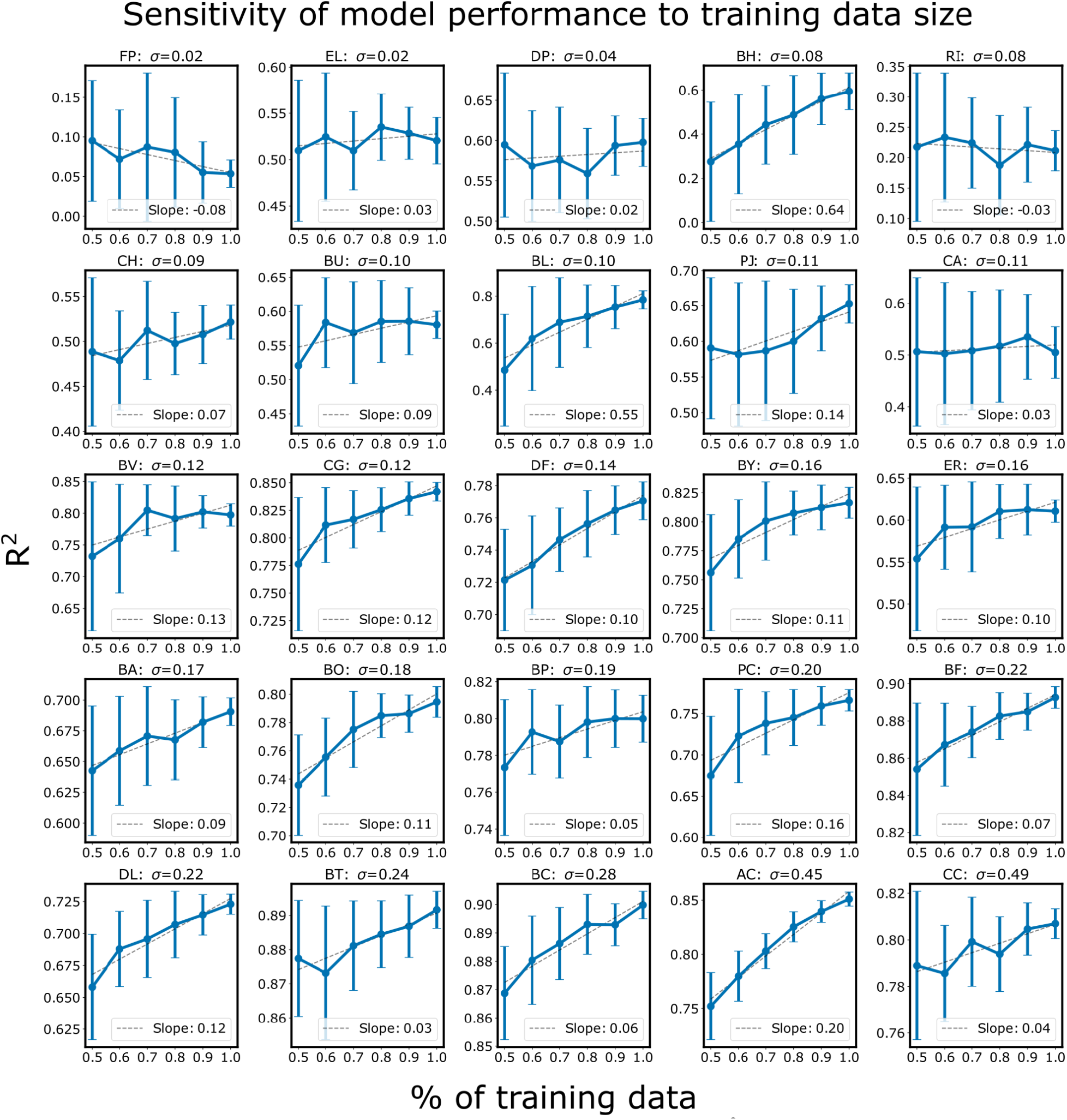
Sensitivity of species abundance prediction performance (R^2^) to the size of the training dataset for each species. Training datasets were randomly subsampled 30 times using 50% to 100% of the total dataset in increments of 10%. Each subsampled training set was subject to 20-fold cross-validation to assess prediction performance. Sub-plots show the mean prediction performance (error bars indicate one standard deviation) over the 30 trials for each percentage of the dataset. Subplots were sorted according to the variance in species abundance taken over the total dataset. In general, prediction performance of low variance species was less likely to improve in response to more training data. N and p-values are reported in **Table S3**.

**Figure S6.**
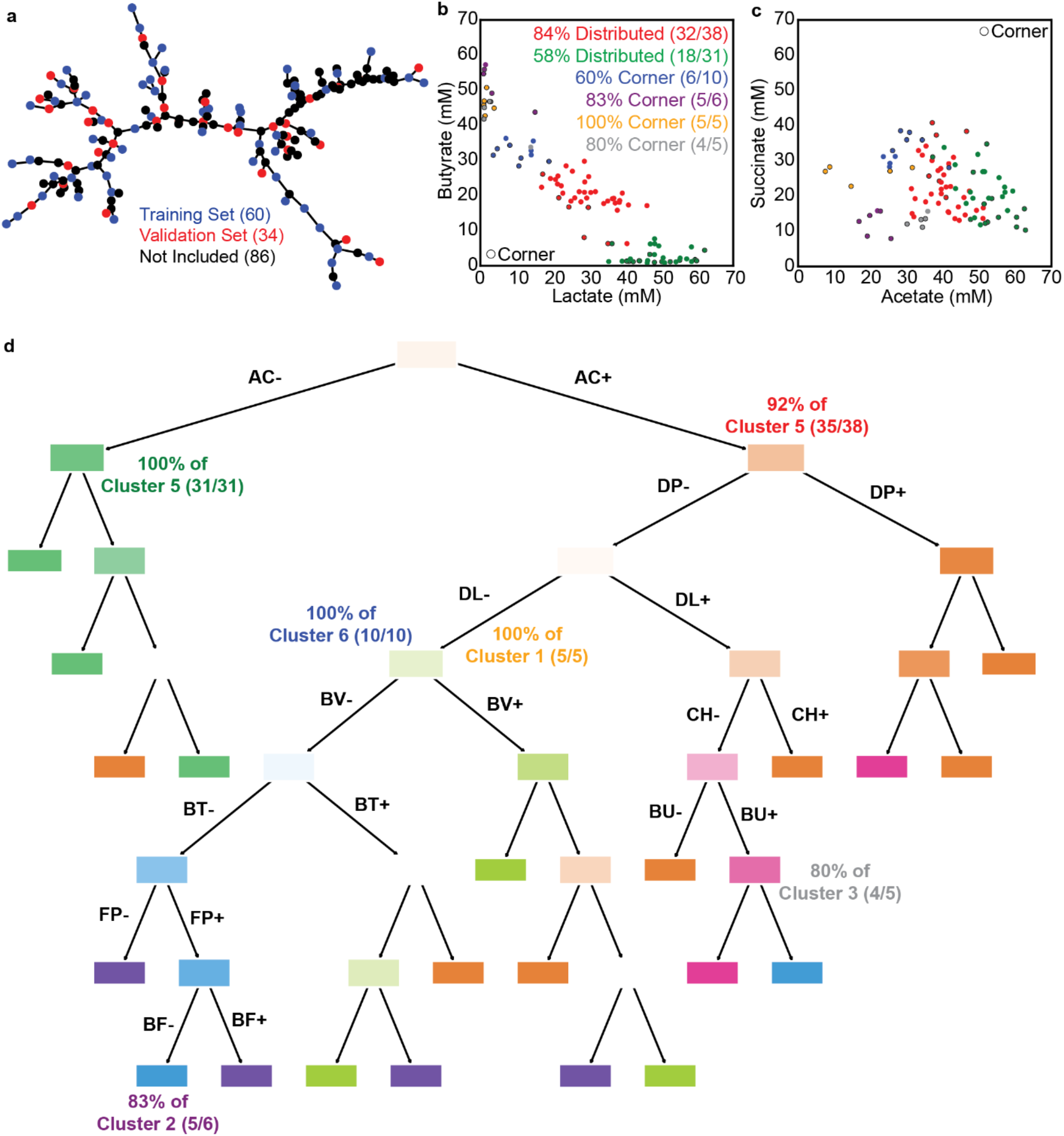
Characteristics of the dynamic community behaviors. (**a**) Minimal spanning tree of a graph representation of the 180 communities characterized in Figure 3 where each node represents a community the lengths of the edges connecting the nodes represent the Euclidean distance between a pair of communities in the 4-dimensional metabolite space. The even spread of the training and validation sets across this tree demonstrate that the subset of communities characterized in the dynamic experiment was representative of all 180 communities characterized in Figure 3. Blue and red nodes indicate the subset of communities chosen for dynamic characterization and used as training and validation examples for LSTM model M3 in **Figure 5**. These subsets were chosen by first performing k-means clustering with *k*=94 for the 180 communities and identifying the 94 communities closest to each cluster centroid and then repeating this process to subsample 34 of the 94 communities (as the training/validation split). (**b**) and (**c**) Scatter plots showing where the clusters from **Figure 5a** fall in the 48-hour metabolite measurement space for comparison with **Figure 3b**. Each datapoint represents a community with the color corresponding to the clusters in **Figure 5a**. Datapoints with black borders come from the “corner” set. Legend indicates the percentage of communities from each cluster that come from the “corner” or “distributed” sets. (**d**) Decision tree classifier explaining which species’ presence determines the clusters of dynamic community behavior from **Figure 5**. Annotations indicate the percentage of communities from each cluster that can be explained by the indicated paths, which are also annotated on **Figure 5a**.

**Figure S7.**
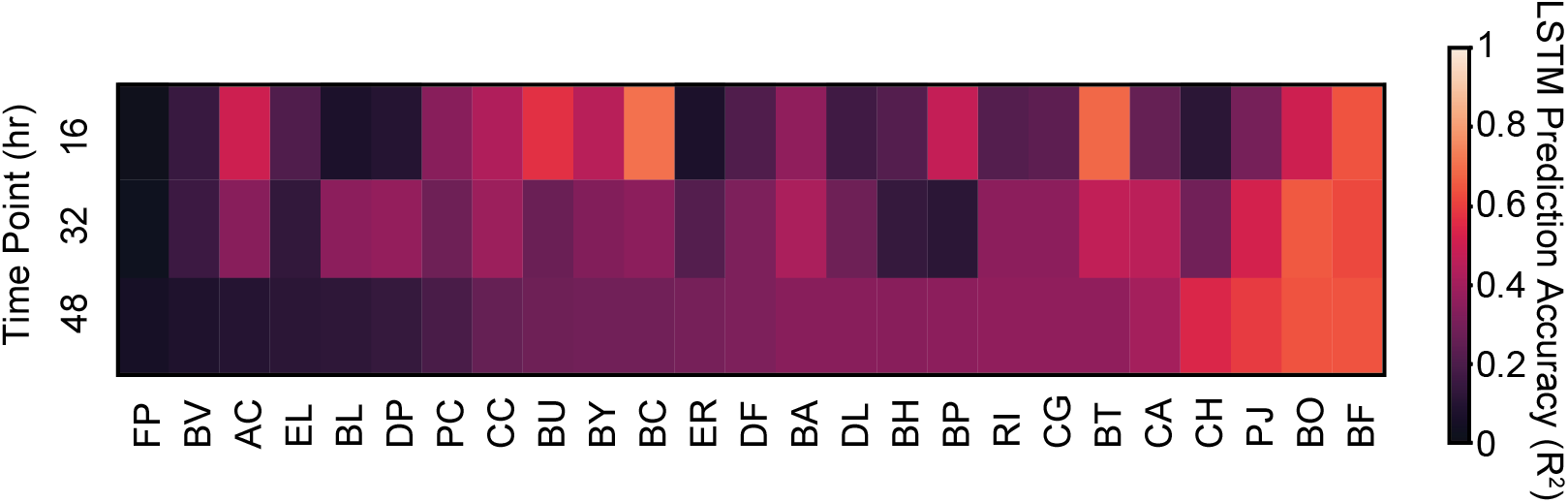
Prediction accuracy of model M3 for species abundance. Heatmap represents R^2^ for the prediction accuracy of model M3 of the abundance of each species at each time point in the 34 validation communities.

